# The impacts of biofuel crops on local biodiversity: a global synthesis

**DOI:** 10.1101/2020.12.21.422503

**Authors:** Sophie Jane Tudge, Andy Purvis, Adriana De Palma

## Abstract

Concerns about the environmental impacts of climate change have led to increased targets for biofuel in the global energy market. First-generation biofuel crops contain oil, sugar or starch and are usually also grown for food, whereas second-generation biofuel is derived from non-food sources, including lignocellulosic crops, fast-growing trees, crop residues and waste. Increasing biofuel production drives land-use change, a major cause of biodiversity loss, but there is limited knowledge of how different first- and second-generation biofuel crops affect local biodiversity. A more detailed understanding could support better decisions about the net environmental impacts of biofuels. We synthesised data from 116 sources where a potential biofuel crop was grown and estimated how two measures of local biodiversity, species richness and total abundance, responded to different crops. Local species richness and abundance were 37% and 49% lower at sites planted with first-generation biofuel crops than in sites with primary vegetation. Soybean, wheat, maize and oil palm had the worst effects; the worst affected regions were Asia and Central and South America; and plant species richness and vertebrate abundance were the worst affected biodiversity measures. Second-generation biofuels had significantly smaller effects: species richness and abundance were 19% and 25%, respectively, lower in such sites than in primary vegetation. Our models suggest that land clearance to generate biofuel results in negative impacts on local biodiversity. However, the geographic and taxonomic variation in effects, and the variation in yields among different crops, are all relevant for making the most sustainable land-use decisions.

## Introduction

Global biodiversity is continuing to decline, with an increasing number of species at risk of extinction (Tittensor et al. 2014; Díaz et al. 2019). Land-use change is currently the biggest threat to biodiversity, followed by overexploitation, climate change and invasive species, and these threats are showing increasing trends (Butchart et al. 2010; Díaz et al. 2019). Land-use change is driven predominantly by agricultural expansion and intensification, and agricultural activities now occupy over 40% of the terrestrial surface (Matson et al. 1997; Foley et al. 2005). Demand for agricultural products is likely to rise further, given the projected increase in human population, heightening the pressure on land, farming and biodiversity in the future (Godfray et al. 2010; United Nations 2019).

While land-use change is currently the major direct threat to terrestrial biodiversity (Díaz et al. 2019), it needs to be balanced with other environmental concerns, including climate change (Thomas et al. 2004). One way of tackling climate change is by increasing the proportion of cleaner, renewable energy sources in the global energy mix (International Energy Agency 2014). Biofuels (derived from plant material) are one such renewable energy source, and are considered a good alternative to fossil fuels due to their lower carbon emissions (Thuiller 2007), although emissions can actually be higher if land is deforested in order to grow biofuel crops (Danielsen et al. 2009; Tilman et al. 2009; Dornburg et al. 2010). Private investment in biofuels has been encouraged by governments worldwide, and by 2012 biofuels met 10% of global energy demand; this demand is likely to triple by 2040 (International Energy Agency 2014), growing particularly quickly in developing countries (Gadonneix et al. 2010).

The market for biofuels consists mainly of bioethanol and biodiesel. These can be produced from food or non-food crops, raising additional concerns about competition with food production (Immerzeel et al. 2014). Biofuels derived from food crops and vegetable oils are classified as first-generation (Correa et al. 2017), such as maize and palm oil. First-generation crops make up most of the global biofuel supply, and are also some of the most intensively farmed crops worldwide, associated with large-scale environmental destruction (Dornburg et al. 2010; Correa et al. 2017). Second-generation biofuels are mainly derived from perennial crops that are not grown for food, including lignocellulosic crops and fast-growing trees, which have lower yields (Dauber et al. 2010). Second-generation biofuels can also be derived from the non-edible parts of food crops (Szymanska-Chargot et al. 2017), from forestry waste and from municipal waste (Koh and Ghazoul 2008).

Palm oil is among the best-known first-generation biofuels. It is the main source of vegetable oil and a major source of biodiesel (Fitzherbert et al. 2008), but expansion of oil palm plantations has already led to large-scale land clearance and deforestation, particularly in Southeast Asia (Dornburg et al. 2010; Barnes et al. 2014). In Malaysia and Indonesia, over half of the expansion between 1990 and 2005 occurred at the expense of forest habitat (Koh and Ghazoul 2008), resulting in significant ecological changes. The uniform structure of plantations compared to natural forest reduces the species diversity and leads to the dominance of generalist species at the expense of specialists, which tend to be of greater conservation concern (Fitzherbert et al. 2008; Phillips et al. 2017). Reduced species richness in oil palm plantations has been reported for a variety of taxa, including birds (Aratrakorn et al. 2006; Peh et al. 2006), bats (Freudmann et al. 2015), invertebrates (Barnes et al. 2014), plants and lizards (Danielsen et al. 2009).

Whereas palm oil is the dominant biofuel crop in Southeast Asia, the USA and Brazil together produce around 80% of global biofuel, in the form of bioethanol, from maize and sugarcane respectively (Gadonneix et al. 2010). Row crops such as maize are more efficient when grown in large monocultures (Wiens et al. 2011), accompanied by the removal of natural features such as copses to produce landscapes with reduced compositional and configurational heterogeneity (Fahrig et al. 2011). The modified landscape affects the availability of resources and the movement and abundance of species (Flather and Bevers 2002; Fahrig et al. 2011). Research into row crops, and other first-generation biofuel crops including oil palm, has found particular declines in vertebrates (Fitzherbert et al. 2008; Fletcher et al. 2011). For example, bird diversity and mammal abundance were found to be over 50% lower in row crop fields in the USA compared to non-crop areas (Fletcher et al. 2011). Intensively managed croplands in the USA and Europe also affect pollinator communities, leading to a reduced species richness and abundance of bees (Steffan-Dewenter et al. 2002; Neumann and Carreck 2010; Kennedy et al. 2013).

Because first-generation biofuels are generally higher-yielding than second-generation biofuels, they are thought to be more damaging to biodiversity, per unit area (Immerzeel et al. 2014; Núñez-Regueiro et al. 2020), and some studies have reported that second-generation biofuels can have positive effects in temperate regions (Robertson et al. 2011; Immerzeel et al. 2014; Haughton et al. 2016). These benefits may relate to several factors, including reduced chemical inputs, longer rotation periods and greater spatial heterogeneity (Dauber et al. 2010). For example, biodiversity benefits have been found for second-generation woody and herbaceous crops including poplar (Christian et al. 1997; Hanowski et al. 1997), willow (Haughton et al. 2016) and perennial grasses (Robertson et al. 2011; Kline et al. 2015; Haughton et al. 2016) when compared to annual row crops. Second-generation crops can also be grown on a wider range of land types including marginal or degraded land, which would not otherwise be suitable for food crops (Wiens et al. 2011), thus providing bioenergy without the need for further land clearance (Immerzeel et al. 2014) or additional competition with food production (Erb et al. 2012).

Previous studies that have considered regional differences have shown that Asia’s biodiversity is affected more strongly by land-use change (Gibson et al. 2011) and conversion to plantations (Phillips et al. 2017) than other regions. The reasons probably relate to the intrinsic sensitivities of the species present and differences in the sampling techniques used, crops grown and local management practices (Gibson et al. 2011; Phillips et al. 2017). Existing comparisons between oil palm plantations (predominant in Asia) and other crops, do indeed highlight oil palm’s particularly detrimental effects. For example, analyses have found lower species richness in oil palm plantations than in second-generation wood, fruit, vegetable, coffee, cocoa and rubber plantations (Peh et al. 2006; Fitzherbert et al. 2008; Barnes et al. 2014; Phillips et al. 2017).

With such a diverse range of crops available for biofuel production and more land continuously being converted to cropland, it is increasingly important to be able to predict how ecological communities will be affected by the different biofuels and to integrate this knowledge into sustainability assessments. However, most studies on biofuels and biodiversity are limited in taxonomic or geographic scope, and there is also a lack of comparisons between crops and between tropical and temperate regions (Dauber et al. 2010; Dornburg et al. 2010; Dauber and Miyake 2016). In this paper, we have made use of the interrelatedness between food and biofuel crops to analyse the biodiversity response to crops using data from published studies that have been collated within the PREDICTS database (Projecting Responses of Ecological Diversity In Changing Terrestrial Systems). The database provides a framework for modelling how site-level biodiversity responds to different land-uses and related pressures, and contains information on the biodiversity in croplands around the world (Hudson et al. 2014; Hudson et al. 2017). We focused our analyses on the biodiversity impacts of first- and second-generation biofuels and compared the effects of 95 biofuel crops in different geographic regions and on different taxonomic groups.

## Methods

### Identifying biofuels in the PREDICTS database

The PREDICTS database (Hudson et al. 2017) contains data from published articles, or reports using published methods, on biodiversity in sites with contrasting land-uses and human pressures around the world. Data for our research was added to the database by the project team between March 2012 and March 2018, following extensive literature searches. Within the database, each source comprises one or more studies (each with a different sampling methodology), which may be arranged in spatial blocks, and that each have data from two or more sites where species richness, abundance or occurrence have been measured using the same procedure, with detail on sampling effort and geographic coordinates (see Hudson et al. 2014 for details).

Each site within the database is assigned a land-use, which can be cropland, pasture, plantation forest, primary forest or non-forest, secondary vegetation (of different ages) or urban (Hudson et al. 2014). Sites are also assigned a land-use intensity – minimal, light or intense use – based on the authors’ descriptions of the sites (Hudson et al. 2014). For our analyses we combined all primary sites into one class and all secondary vegetation sites into another, and extracted from the database the sites where the predominant habitat was cropland or plantation forest and where the name of the crop grown was known, using site-specific crop data from Hill et al. (2018). We then conducted a literature search of the crops to find out which can be used as biofuels, using the scientific or common crop name “AND” a variety of biofuel terms, including biofuel, biodiesel, bioethanol, ethanol, fuel, energy and bioenergy. When the crop name was too broad, such as the family name Poaceae, it was excluded from the search, as it was not possible to tell which species or subspecies were grown at the site. However, some genus names were included when it was still possible to assess their suitability, such as wheat. We classified each biofuel crop as first- or second-generation (Table 1) and into categories of similar crops. First-generation biofuel crops had the highest proportion of sites where land-use was classified as intensely managed within the database, rather than light, minimal or unknown intensity of use (48% for first-generation crops compared to 16% for second-generation crops) (Table 2). Because our main focus is the differences between, rather than within, biofuel generations and categories, we did not use land-use intensity as an explanatory variable, but interpreted our results assuming that sites with first-generation biofuels tend to be more intensively managed than sites with second-generation biofuels.

**Table 1.** Criteria for sites within the PREDICTS database to be classed as first-generation biofuel or second-generation biofuel for our analyses.

| Biofuel generation | Inclusion criteria |
| --- | --- |
| First | Site has a single first-generation biofuel crop<br>Site has more than one biofuel crop, all of which are first-generation |
| Second | Site has a single second-generation biofuel crop<br>Site has more than one biofuel crop, all of which are second-generation |

**Table 2.** Land-use intensity for each site from the PREDICTS database that was used for our analyses. See Hudson et al. (2014) for land-use intensity descriptions.

| Land-use | Number of sites with each land-use intensity |  |  |  |
| --- | --- | --- | --- | --- |
|  | Cannot decide | Intense use | Light use | Minimal use |
| First-generation<br>biofuel | 41 | 258 | 220 | 24 |
| Second-generation<br>biofuel | 106 | 138 | 335 | 282 |
| Primary vegetation | 464 | 617 | 2556 | 5204 |
| Secondary<br>vegetation | 1452 | 723 | 1804 | 3196 |
| Pasture | 1091 | 1476 | 2305 | 1609 |
| Urban | 41 | 287 | 756 | 401 |

Very few sources explicitly stated whether crops at each site were being grown for biofuel. We therefore treated all sites with biofuel crops as biofuel sites, as in Fletcher et al. (2011) and Núñez-Regueiro et al. (2020), assuming that the biodiversity effects do not depend on the final use of the crop. Analysis of data from sites that are actually being used for biofuel would be preferable and more informative.

### Statistical modelling

We modelled the response of biodiversity to biofuel crop category and biofuel crop generation, comparing the results with the other PREDICTS land-uses (primary vegetation, secondary vegetation, pasture and urban) (Hudson et al. 2014). As measures of local biodiversity, we used species richness and total abundance. Although other biodiversity measures that reflect species composition and the relative abundances of the species present contain more information and are more sensitive to biodiversity change (Santini et al. 2016; Hillebrand et al. 2018), most papers only report a single measure of biodiversity, most commonly species richness (Naeem et al. 2016). We tested whether the effects of first- and second-generation biofuel crops differed significantly among geographic regions (Africa, Asia, Europe, Central and South America, North America and Oceania) and major taxonomic groupings (plants, vertebrates and invertebrates). Thus, in all, we fitted eight models: two to describe the overall effects of biofuel crop generation on species richness and total abundance, two for the effects of biofuel crop category on species richness and total abundance, two for the effects of biofuel crop generation in different regions on species richness and total abundance, and two for the effects of biofuel crop generation on the species richness and total abundance of different taxonomic groups. These were all fitted as linear mixed-effects models using the “lme4” package (Bates et al. 2015), as such models are suitable when dealing with a nested data structure (e.g. there are differences in sampling methods and sampling effort between studies) (Phillips et al. 2017; Núñez-Regueiro et al. 2020). The mixed-effects models were able to account for the non-independence of biodiversity measures within studies (SS: source-study) and blocks (SSB: source-study-block) by fitting them as random intercepts.

Species richness models were fitted with a Poisson error structure and a log link function (Zuur et al. 2009); to deal with overdispersion otherwise seen in these models, a site level random intercept was included (SSBS: source-study-block-site) (Newbold et al. 2015). Total abundance is not always an integer, so a Poisson error structure could not be used; instead, data were log(x+1) transformed and modelled with a Gaussian error structure. For each of the eight models, we first compared different random intercept structures and chose the structure with the lowest Akaike Information Criterion. We did not include random slopes to increase the chance of model convergence. To choose the best fixed-effects structure we used backwards stepwise model simplification of fixed effects using likelihood ratio tests (Zuur et al. 2009). The fixed effects that were considered included the interaction between biofuel generation and region and between biofuel generation and taxon, but we did not include three-way interactions.

When a model failed to converge, we increased the number of iterations and then used the “allFit()” function from the “lme4” package to compare the estimated values from all available different optimizers. If all optimizers gave very similar estimates (within 0.01), we considered the converge warnings to be false positives (Bates et al. 2015). We also used the “MCMCglmm” package and compared the outputs, verifying that values were within 0.01 of each other (Hadfield 2010).

For the final models, the “MuMIn” package was used to calculate the marginal R^2^ (amount of variance explained by fixed factors) and the conditional R^2^ (amount of variance explained by fixed and random factors) (Bartoń 2018). Additionally, the “car” package was used to conduct type II anova tests on the models (Fox and Weisberg 2017). All analyses were carried out using R version 3.4.2 (R Core Team 2017).

## Results

### Biofuels in the PREDICTS database

At the time of our analyses, the PREDICTS database contained data from 32,076 sites and 552 sources. At least one crop name was available from 4,033 sites (159 sources), giving a total of 150 names, although some were different spellings of the same crop. Excluding those where the names were too vague, we assessed 95 unique crops for biofuel potential. Of the 95 crops, we found research identifying 65% (62 crops) as a biofuel by at least one source, with the majority (49) of the crops having waste that could be used for biofuel. Supplementary Material 1 gives detail of our assessment of the crops’ biofuel potential and grouping into biofuel generations and categories. Of the biofuel crops identified, 26 were first-generation and 36 were second-generation. We grouped the biofuel sites into the following categories, to enable robust statistical modelling: coffee, cotton, fruit/vegetable, maize, mixed crops, oil palm, other grain, other oil crop, perennial grass, rapeseed oil, rubber, soybean and wheat. There were not enough data (only three sites) to include woody crops in our analysis of the effects of biofuel category on biodiversity. Our dataset included 543 first-generation biofuel sites and 861 second-generation biofuel sites. The geographic spread of data was uneven, with the majority of first-generation sites being in Europe and second-generation sites in Central and South America. Due to imbalance in the variety of crops grown in each region and the taxonomic groups recorded in each crop category, we did not model the effects of individual crop categories in different regions or on different taxonomic groups separately (see Supplementary Material 2).

### Biofuels and biodiversity

Biofuel crop generation had a significant effect on the total abundance (χ^2^=118.21, df=5, p<0.001) and species richness (χ ^2^=443.39, df=5, p<0.001) of sites. Overall, species richness was 37% lower (estimate=-0.45, SE=0.03, p<0.001; Figure 1) and total abundance 39% lower (estimate=-0.49, SE=0.07, t=-7.15; Figure 2) in first-generation biofuel sites than in primary vegetation, which were the biggest declines of all the land-uses assessed. Second-generation biofuel sites had a total abundance 25% (estimate=-0.29, SE=0.06, t=-5.2) lower than primary vegetation, which was lower than pasture, secondary vegetation and urban sites.

**Figure 1.**
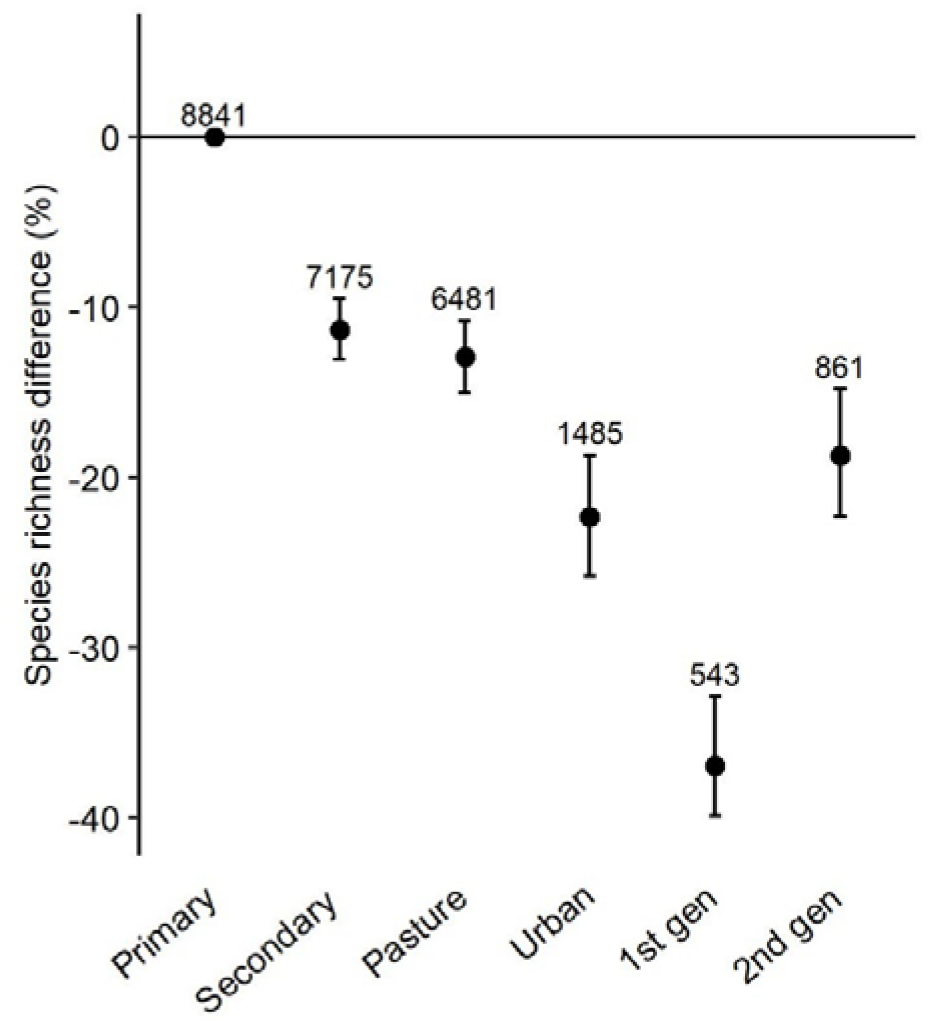
Species richness in sites with first- and second-generation biofuel crops and reference land-uses from the PREDICTS database: primary vegetation, secondary vegetation, pasture and urban. Error bars show 95% confidence intervals. Error bars that do not cross zero are considered significantly different to the baseline, primary vegetation. Data point labels show sample size i.e. number of sites.

**Figure 2.**
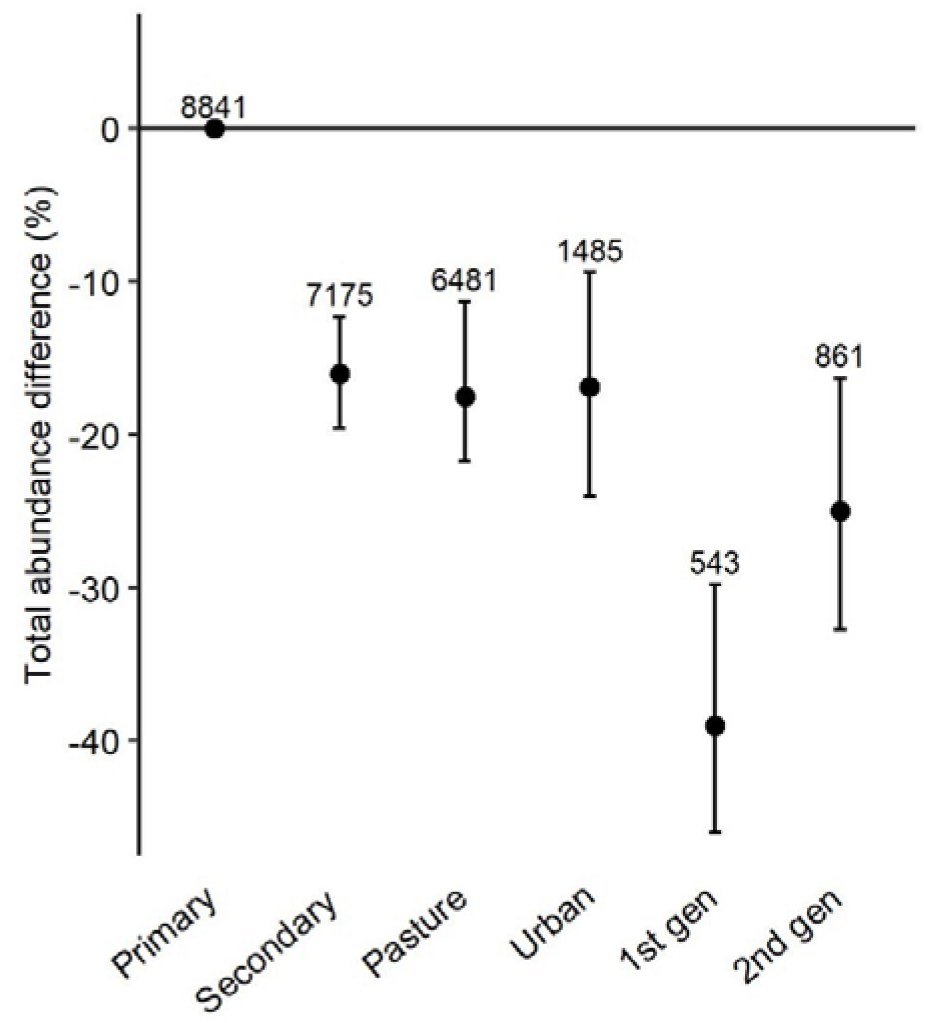
Total abundance in sites with first- and second-generation biofuel crops and reference land-uses from the PREDICTS database: primary vegetation, secondary vegetation, pasture and urban. Error bars show 95% confidence intervals. Error bars that do not cross zero are considered significantly different to the baseline, primary vegetation. Data point labels show sample size i.e. number of sites.

The type of biofuel crop (category) also had a significant effect on total abundance (χ ^2^=313.21, df= 16, p<0.001) and species richness (χ ^2^=518.32, df=16, p<0.001). Sites where cotton and soybean were grown recorded the lowest species richness, followed by wheat and maize (Figure 3). Species richness in sites with other grain crops and rubber was not significantly different from that in primary vegetation (although the confidence intervals were wide), whereas all other crop categories had a significantly lower species richness. Sites with cotton also had a very low total abundance of organisms (86% less than primary vegetation), followed by soybean and oil palm. On the other hand, abundance in sites planted with rubber, other oil crops and fruit/vegetable did not differ significantly from that in primary vegetation, and perennial grass and rapeseed oil crops had total abundances more than 50% higher (Figure 4).

**Figure 3.**
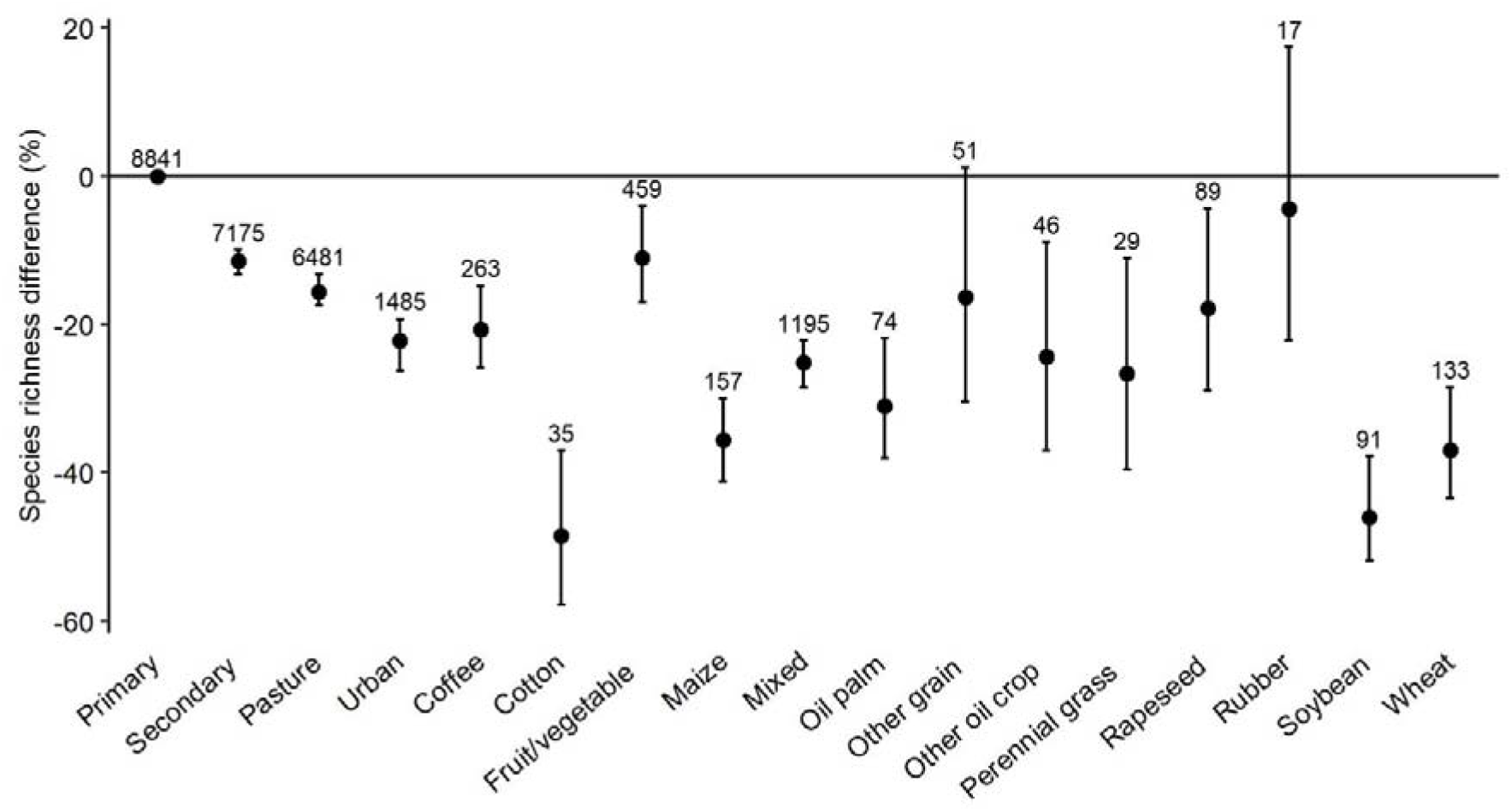
Species richness in biofuel crop categories and reference land-uses from the PREDICTS database: primary vegetation, secondary vegetation, pasture and urban. Error bars show 95% confidence intervals. Error bars that do not cross zero are considered significantly different to the baseline, primary vegetation. Data point labels show sample size i.e. number of sites.

**Figure 4.**
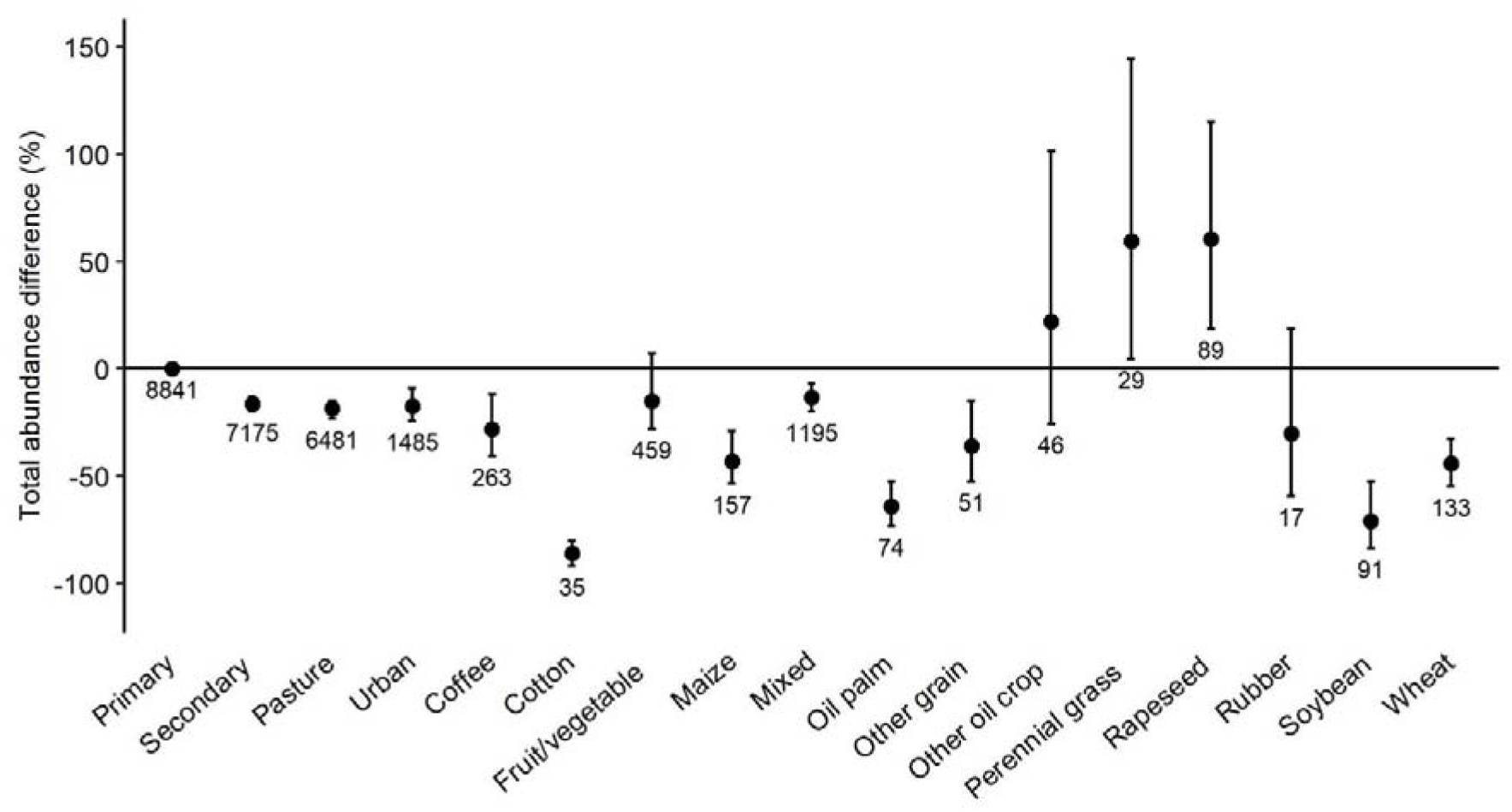
Total abundance in biofuel crop categories and reference land-uses from the PREDICTS database: primary vegetation, secondary vegetation, pasture and urban. Error bars show 95% confidence intervals. Error bars that do not cross zero are considered significantly different to the baseline, primary vegetation. Data point labels show sample size i.e. number of sites.

### Geographic variation

The effects of the two generations of biofuel crops on biodiversity varied significantly among regions (Table 3). From our data, sites with first-generation biofuels supported fewer species on average than sites with second-generation biofuels for all regions (Figure 5). Within the first-generation group, all regions had significantly lower species richness than primary vegetation (apart from North America, which showed wide variability), between 40% lower in Asia and 31% lower in Europe. In our model of croplands with second-generation biofuels, species richness was significantly lower than primary vegetation only in Africa, Asia and Central and South America.

**Figure 5.**
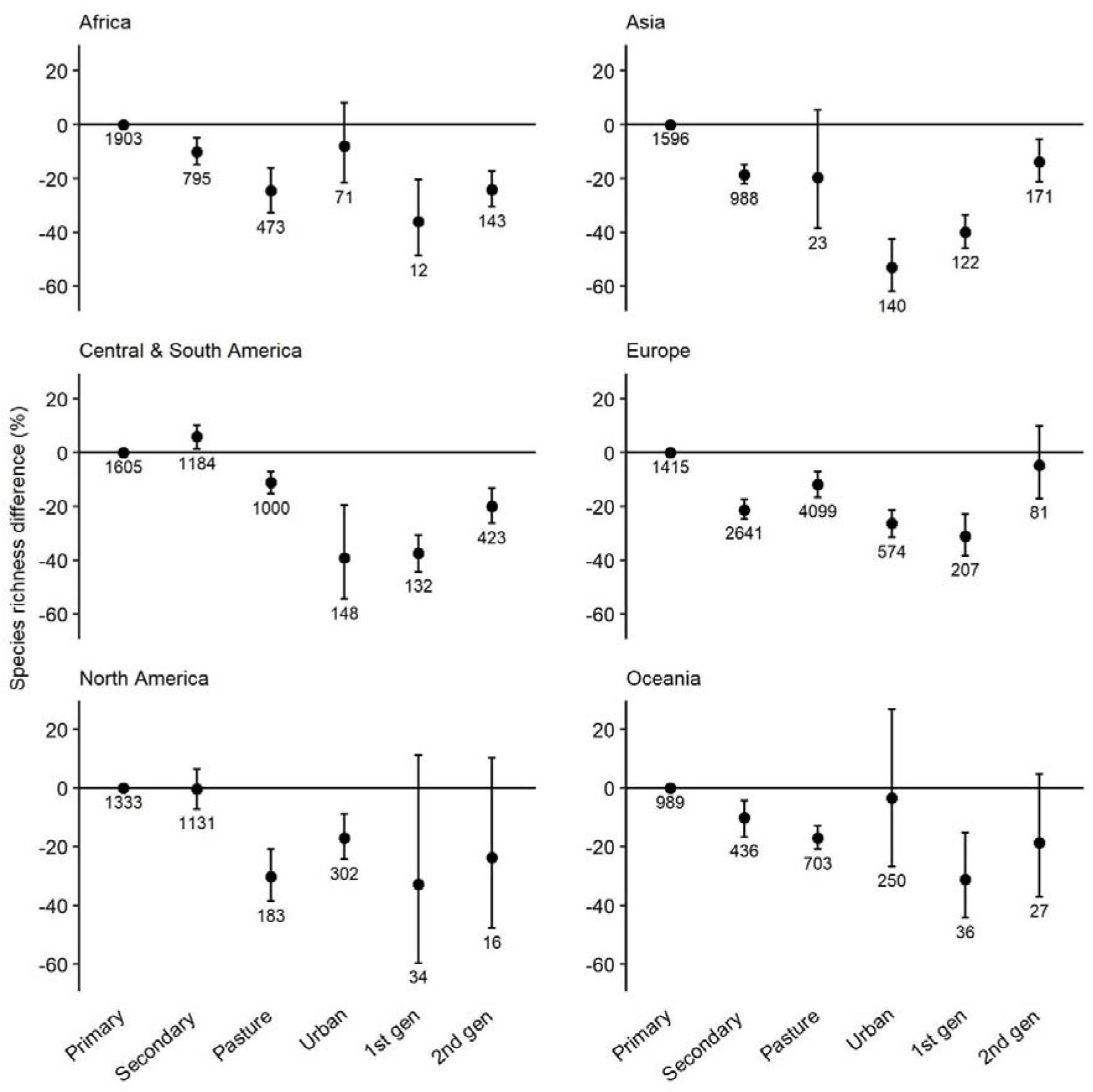
Species richness in sites with first- and second-generation biofuel crops and reference land-uses from the PREDICTS database: primary vegetation, secondary vegetation, pasture and urban, in Africa, Asia, Central and South America, Europe, North America and Oceania. Error bars show 95% confidence intervals. Error bars that do not cross zero are considered significantly different to the baseline, primary vegetation. Data point labels show sample size i.e. number of sites.

**Table 3.**
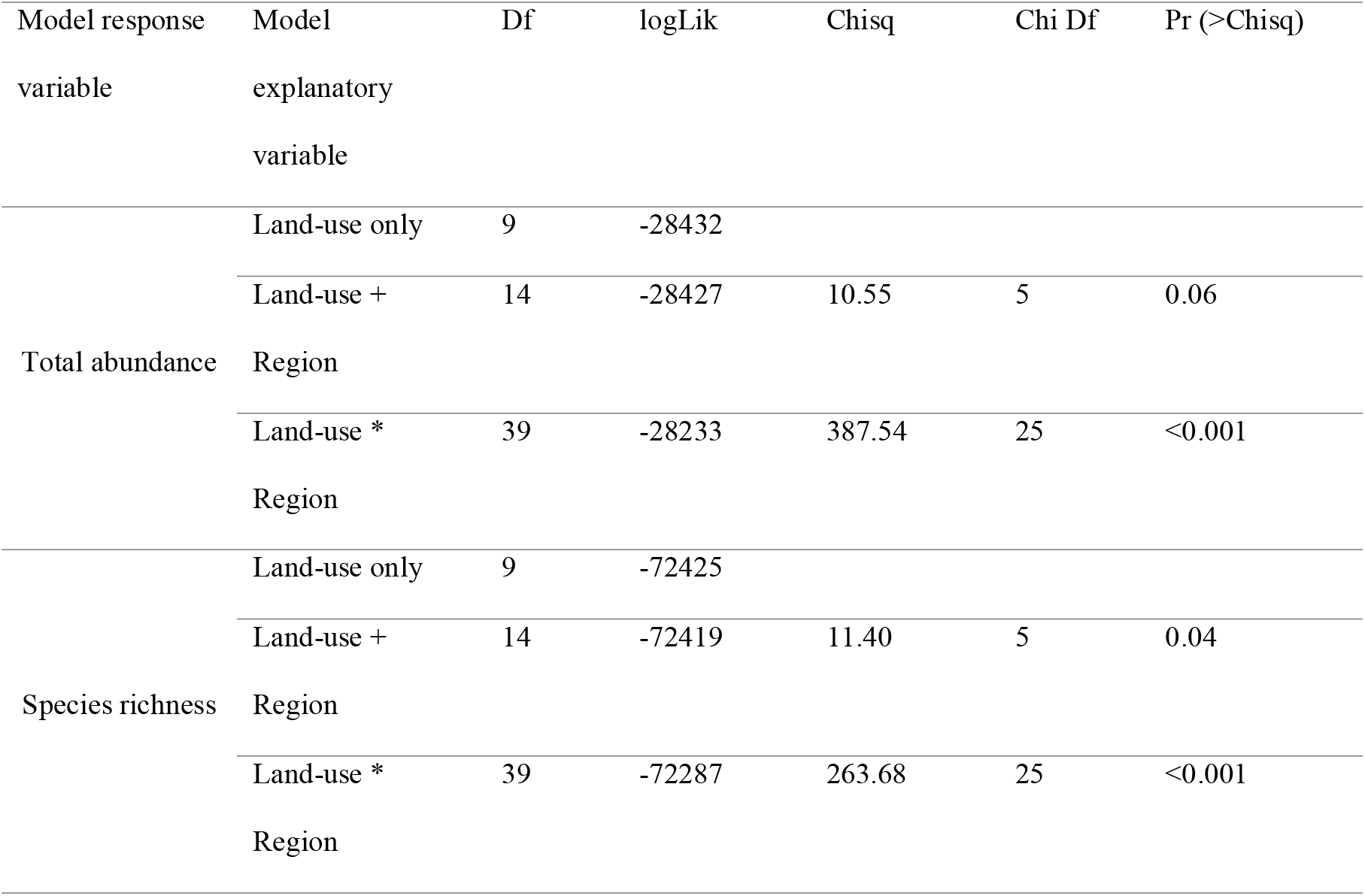
Type II anova results for minimum adequate models with total abundance and species richness as the response variables and land-use (including biofuel generation), region and their interaction as the explanatory variables. Chisq is the result of a chi-square test, Df is degrees of freedom and logLik is log-likelihood. Pr (>Chisq) is a measure of the significance of the model explanatory variable shown.

First-generation biofuel sites also had a lower total abundance of organisms than second-generation sites in all regions except for Africa, where abundance was lowest in second-generation sites. First-generation biofuels grown in Central and South America and Asia had the biggest impact on total abundance across all the regions, according to our model (Figure 6). In Europe, all land-uses supported lower total abundance than primary vegetation. On the other hand, in North America only pasture had a lower total abundance than primary vegetation, and second-generation biofuels increased it by 138% (however, there were very large confidence intervals). Similarly, in Oceania, we found that biofuels had no significant effect on total abundance.

**Figure 6.**
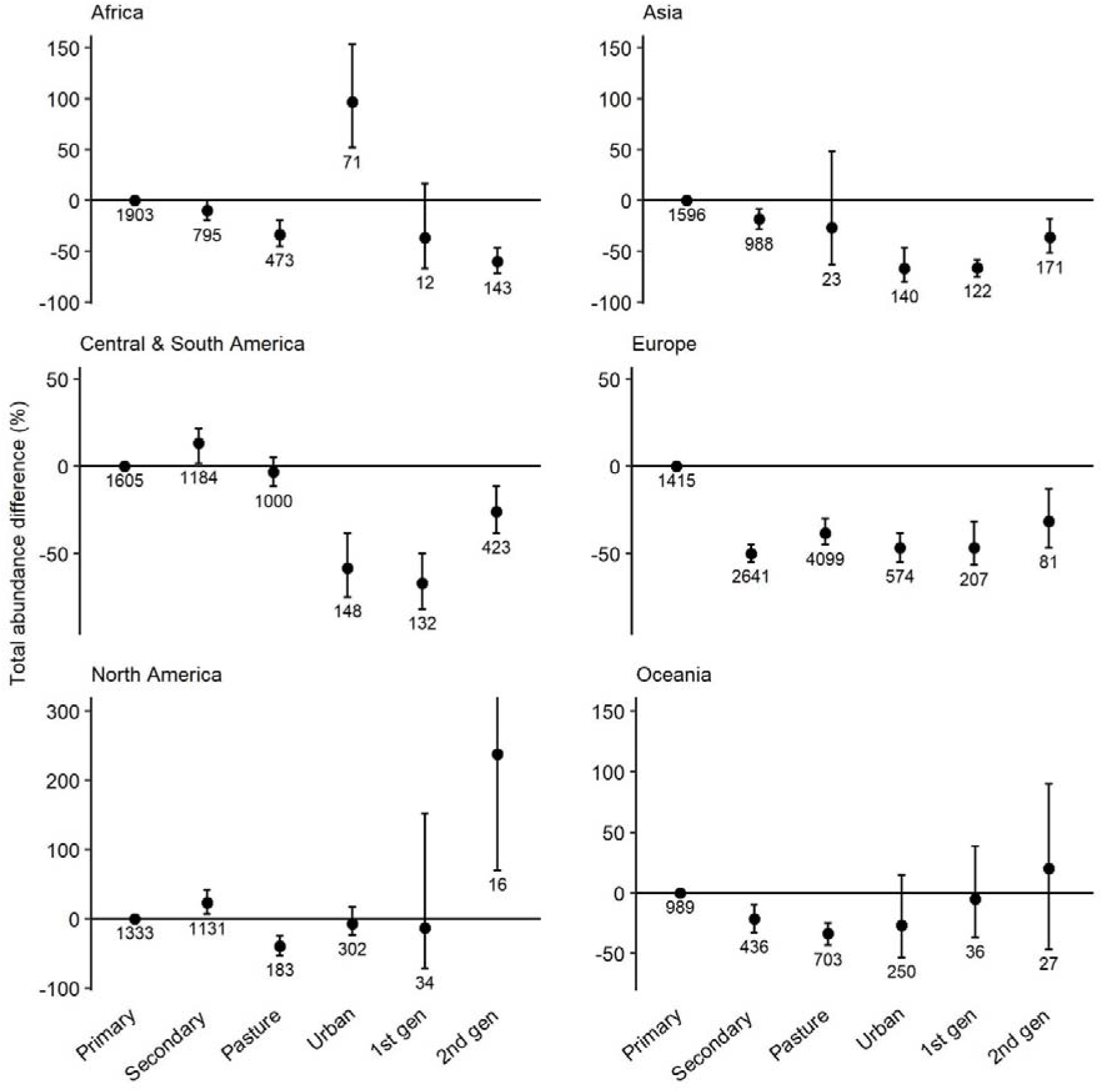
Total abundance in sites with first- and second-generation biofuel crops and reference land-uses from the PREDICTS database: primary vegetation, secondary vegetation, pasture and urban, in Africa, Asia, Central and South America, Europe, North America and Oceania. Error bars show 95% confidence intervals. Error bars that do not cross zero are considered significantly different to the baseline, primary vegetation. Data point labels show sample size i.e. number of sites.

### Taxonomic variation

The interaction between taxonomic group and biofuel generation was significant for both species richness and total abundance (Table 4). Croplands with first-generation biofuels had lower species richness of vertebrates (by 28%), invertebrates (by 31%) and plants (by 49%) than did primary vegetation (Figure 7). First-generation sites showed a lower species richness than second-generation sites for all taxa, and for invertebrates and plants this was the lowest value of all the land-uses. The total abundance of plants was not significantly affected by first-generation biofuels, but was 46% lower in second-generation sites (Figure 8). However, the site-level total abundance of vertebrates and invertebrates was significantly lower in both generations of biofuel, and lower in first-than second-generation sites. The worst effect of biofuels, and indeed any land-use, was on the abundance of vertebrates in first-generation sites, which was 69% lower than in primary vegetation. The only positive effect of land-use on biodiversity in these analyses came from the abundance of vertebrates in urban environments.

**Figure 7.**
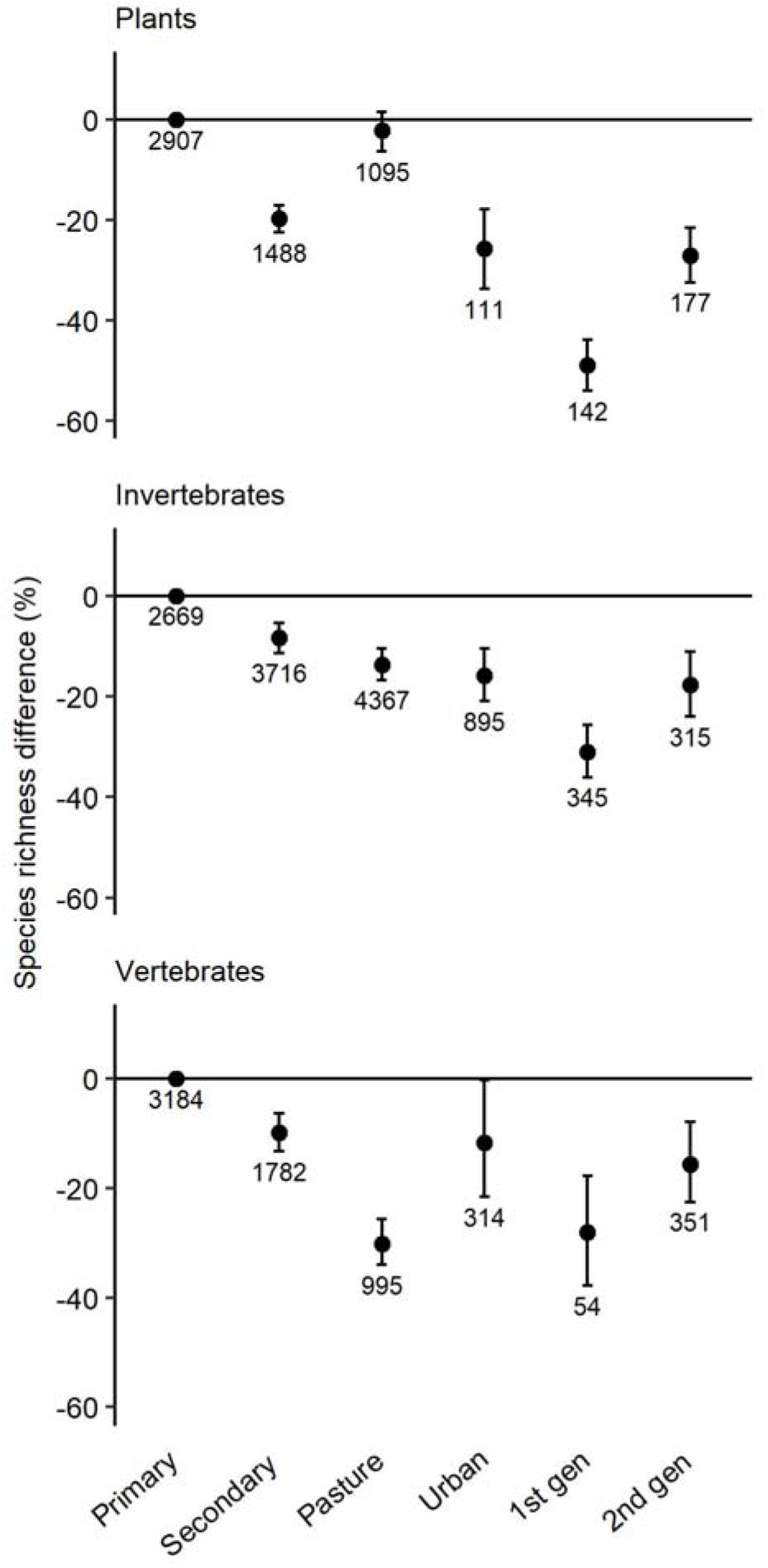
Species richness in sites with first- and second-generation biofuel crops and reference land-uses from the PREDICTS database: primary vegetation, secondary vegetation, pasture and urban, for plants, invertebrates and vertebrates. Error bars show 95% confidence intervals. Error bars that do not cross zero are considered significantly different to the baseline, primary vegetation. Data point labels show sample size i.e. number of sites.

**Figure 8.**
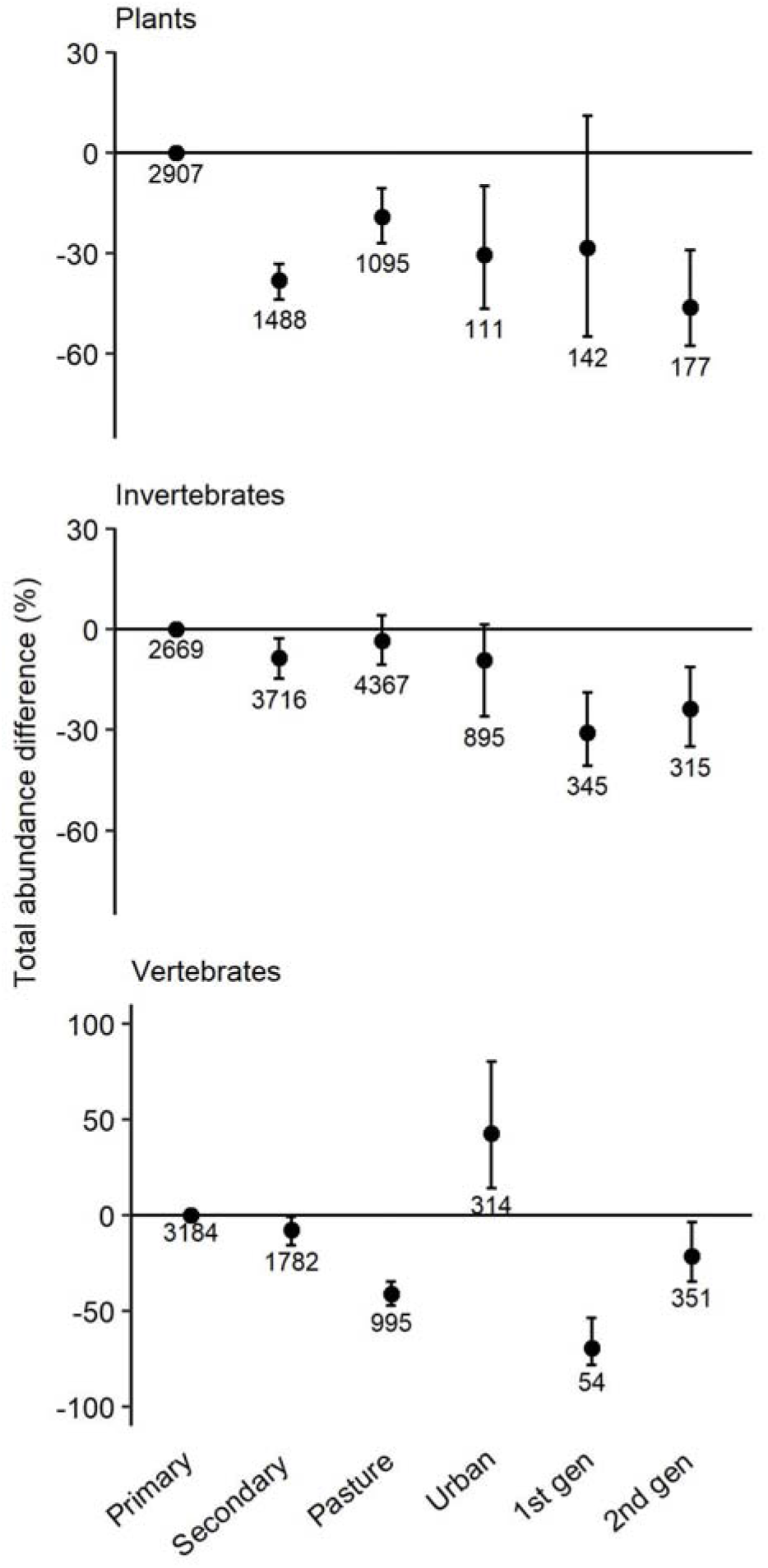
Total abundance in sites with first- and second-generation biofuel crops and reference land-uses from the PREDICTS database: primary vegetation, secondary vegetation, pasture and urban, for plants, invertebrates and vertebrates. Error bars show 95% confidence intervals. Error bars that do not cross zero are considered significantly different to the baseline, primary vegetation. Data point labels show sample size i.e. number of sites.

**Table 4.** Type II anova results for minimum adequate models with total abundance and species richness as the response variables and land-use (including biofuel generation), taxon and their interaction as the explanatory variables. Chisq is the result of a chi-square test, Df is degrees of freedom and logLik is log-likelihood. Pr (>Chisq) is a measure of the significance of the model explanatory variable shown.

| Model response variable | Model explanatory variable | Df | logLik | Chisq | Chi Df | Pr (>Chisq) |
| --- | --- | --- | --- | --- | --- | --- |
| Total abundance | Land-use only | 9 | -27979 |  |  |  |
|  | Land-use + Taxon | 11 | -27955 | 47.41 | 2 | <0.001 |
|  | Land-use * Taxon | 21 | -27869 | 173.40 | 10 | <0.001 |
| Species richness | Land-use only | 9 | -70844 |  |  |  |
|  | Land-use + Taxon | 11 | -70829 | 29.69 | 2 | <0.001 |
|  | Land-use * Taxon | 21 | -70736 | 185.75 | 10 | <0.001 |

For the eight final model outputs, see Supplementary Material 3-10.

## Discussion

We conducted a global synthesis exploring the impacts of biofuels on local biodiversity, comparing the largest range of biofuel crops and potential biofuel crops to date (Immerzeel et al. 2014; Núñez-Regueiro et al. 2020). Our results corroborate previous findings that biofuel crops are damaging to biodiversity (Robertson et al. 2011; Immerzeel et al. 2014; Haughton et al. 2016; Núñez-Regueiro et al. 2020), whilst providing new insights into the differences globally and regionally. Our results showed that traditional first-generation biofuel crops including soybean, maize and oil palm were the most detrimental to local species richness and total abundance, worse even than urban environments. Although comparatively better, second-generation biofuel crops were also damaging to these measures of biodiversity overall, but with differences between individual crops. Due to the variation identified, biofuel policies should be tailored to consider how local biodiversity might respond to particular crops, grown in particular geographic regions.

The finding that first-generation biofuels were on average more harmful to local species richness than second-generation biofuels in all regions is not surprising given their higher use-intensity. They tend to be monocultures with significant chemical inputs, no crop rotations, recent clear-felling etc. (Hudson et al. 2014). Many studies have now linked agricultural intensification to a loss of biodiversity, consistent with our results (e.g. Krebs et al. 1999; Robinson and Sutherland 2002; Kennedy et al. 2013). We found that soybean (the most important source of biodiesel in Brazil, one of the world’s largest biodiesel producers (Cerri et al. 2017)) severely reduced local biodiversity, more so than other row crops including maize (Fargione et al. 2010; Poggio et al. 2013). Yet maize and wheat were still very damaging compared to primary and secondary vegetation and second-generation biofuels, reflecting the homogeneity and limited resource availability in row crop landscapes (Smith et al. 2010).

In our models, soybean also supported fewer species than oil palm plantations, whereas Fargione et al. (2010) found the opposite to be true, highlighting differences that can arise depending on data sources and study systems. Nonetheless, our results agree with previous studies that oil palm plantations support fewer species than second-generation coffee, fruit/vegetable and rubber plantations (Peh et al. 2006; Fitzherbert et al. 2008; Barnes et al. 2014; Phillips et al. 2017). Analyses of the greenhouse gas emissions arising from production of these biofuels, using life cycle assessment methods, have had varied results, but a recent analysis by Meijide et al. (2020) found that biofuel generated from young palm oil plantations releases more greenhouse gas emissions than fossil fuels, with consideration given to soil emissions and emissions during the cultivation, milling and fuel production stages. Therefore, theses crops’ environmental benefits are unclear.

Despite second-generation biofuels being better for biodiversity than first-generation biofuels on average, cotton – a second-generation crop – was the most damaging of all the crops analysed. Cotton farming typically involves heavy use of pesticides, which are known to have negative effects on the environment (Carpenter et al 2002). The use of transgenic cotton varieties could alleviate some of the negative impacts (Bouyer et al. 2007), but our results indicate that cotton expansion for biofuel production would not be desirable. Using waste products from existing plantations could remain an option, as for many of the other crops analysed, although care would need to be given to ensure that this does not cause further land clearance. Also, removing crop residues from some plantations could have negative consequences for soil fertility and the diversity of fungi, beetles and birds (Wiens et al. 2011; Ranius et al. 2014), and there are technical challenges to overcome before it can become large-scale (Gadonneix et al. 2010). However, some second-generation crops can be grown on waste or marginal lands that are less suitable for the more profitable crops (Warren-Thomas et al. 2015; Conkling et al. 2017).

Perennial grass – reported as one of the more favourable biofuel crops for biodiversity (Robertson et al. 2011) – and rapeseed oil both had a positive effect on total abundance in our analyses. Perennial grass can provide habitat for migratory birds (Immerzeel et al. 2014) by mimicking natural grassland vegetation structure (Dornburg et al. 2010; Blank et al. 2016), while pollinators can benefit from the presence of flowering rapeseed oil crops (Westphal et al. 2003). However, we found a lower species richness in these croplands, which could have a substantial effect on the ecosystem functioning and conservation value of the land.

Of all the biofuel crop categories in our models, rubber had the most similar species richness to primary vegetation. It also has favourable fuel properties compared to other crops, including soybean (Ikwuagwu et al. 2000). The rubber (and oil palm) sites in our analyses are all in Asia, a region where rubber is expanding rapidly; in 2012 it covered an area of land equivalent to 71% that of oil palm in Southeast Asia (Warren-Thomas et al. 2015). We found a large variability in its effects on biodiversity, which reflects the wider literature. The thin canopy of rubber plantations allows light through, stimulating the growth of understory layers and providing habitat for other species (Peh et al. 2006). The oil-rich seeds may attract small mammals (Nakagawa et al. 2006) and increasing the distance between trees, retaining older trees and using agroforestry techniques can all help to increase the biodiversity value of rubber (Beukema et al. 2007; Mingxia et al. 2017). Our results agree to the extent that vertebrates were less affected by second-generation biofuel crops than invertebrates and plants. Contrastingly, other studies have found a reduced species richness in rubber plantations (Peh et al. 2006; Warren-Thomas et al. 2015; Mingxia et al. 2017). Further research is therefore needed to evaluate the reasons for the variation in response to rubber and to find which management techniques could be used to minimise its ecological impacts.

Asia and Central and South America showed the biggest effect sizes of biofuels, consistent with previous findings that there is geographic variability in the response of biodiversity in disturbed tropical forests and plantations (Gibson et al. 2011; Phillips et al. 2017). The regional differences we found can be partly attributed to variation in the crops grown in each region. For example, Central and South America had the most soybean sites and all oil palm sites were from Asia, both these crops being among the most detrimental first-generation crops. The Central and South America region also grows a lot of sugarcane for biofuel, but our data had no sugarcane sites, and therefore does not show the whole picture (Gadonneix et al. 2010). However, our results highlight the damaging nature of soybean croplands in the region. Regional differences could also be due to inherent differences in the resilience of the ecological communities, especially since most croplands in the tropics are created by replacing tropical forests, which are highly biodiverse ecosystems that are sensitive to disturbance (Phillips et al. 2017).

In North America, there was a large variability in the impacts of biofuels. Most sites had mixed crops or other oil crops. There were only three maize sites in our data for North America, yet maize is widely grown for biofuel in the region (Gadonneix et al. 2010); including more maize sites in our data would probably have led to a more evident negative response to first-generation biofuels. In Africa, all effects in biofuel croplands were negative, though first-generation sites were lacking. A more even and representative spread of crops across regions would have improved our models. In Europe, Oceania and North America, our results showed that second-generation biofuels were not as damaging to biodiversity, highlighting the potential of second-generation biofuels in these regions for creating less environmentally damaging biofuel. However, after rapeseed, soybean and palm oil are still likely to be the greatest sources of biodiesel (contributing 13% each to total biofuel) in Europe, and maize the most important source of bioethanol (Valin et al. 2015). Similarly, in North America, the renewable fuel standard (RFS2) (2010) estimates that by 2022, 42% of renewables in transport fuels will still come from conventional biofuels, derived mainly from first-generation maize starch (Sorda et al. 2010).

When considering the overall responses of different taxa, species richness of plants, invertebrates and vertebrates declined in both first- and second-generation biofuel sites, which is concerning given the projected expansion of the biofuel industry (International Energy Agency 2014; Núñez-Regueiro et al. 2020). For plants, the lower species richness coupled with the limited change in abundance could signal the over-dominance of disturbance-tolerant species at the expense of more specialist species (Phillips et al. 2017), which could lead to biotic homogenisation and the absence of certain species. For example, a meta-analysis by Danielsen et al. (2009) found complete exclusion of epiphytic orchids and indigenous palms in oil palm plantations. Our results also showed that vertebrates were dramatically reduced in abundance in first-generation sites. Data limitations meant we did not test for variability within the vertebrate group. However, Núñez-Regueiro et al. (2020) found a strong increase in the abundance of mammals in ecosystems with biofuel crops, but a strong decrease in the abundance of birds. Additional analyses that consider the compositional similarity of sites with biofuel crops compared to other land-uses and breakdown biodiversity into different indices, species IUCN conservation status, trophic guild and native or alien status would provide useful information for conservation purposes in the future.

Although first-generation biofuels were more damaging to local biodiversity than second-generation biofuels, crop yield can influence which strategy might be best for conserving biodiversity. Lower-yielding second-generation biofuels generally produce less energy per unit area than first-generation biofuels, requiring more land clearance to generate the same amount of fuel, multiplying the negative impacts on biodiversity from habitat loss (Erb et al. 2012). Table 5 compares the energy yield of some of the biofuel crops we analysed, and can be combined with the crops’ effects on local species richness and abundance to compare the cost (in terms of local biodiversity) per unit energy of different crops. For example, although first-generation oil palm’s impact on local species richness is about 1.3 times greater than that of second-generation jatropha, its biofuel yield is at least twice as great (Table 5), meaning that the impact per unit of energy is less with oil palm than with jatropha. Any such conclusions must come with many caveats at present: we have shown that the effects of crops on biodiversity vary among regions and among taxa, and they are likely to also depend on other interacting factors such as farming practices, surrounding habitat and landscape characteristics; and biodiversity measures that incorporate species turnover are more sensitive to anthropogenic change than the measures we have used (Hillebrand et al. 2018). However, this kind of analysis can potentially help decide whether a land-sparing (more intensive, higher-yielding farming on a smaller land area) or a land-sharing (less intensive, lower-yielding farming on a greater land area) approach to future biofuel production would be better for biodiversity and therefore more sustainable (Barbier and Burgess 1997; Phalan et al. 2011).

**Table 5.**
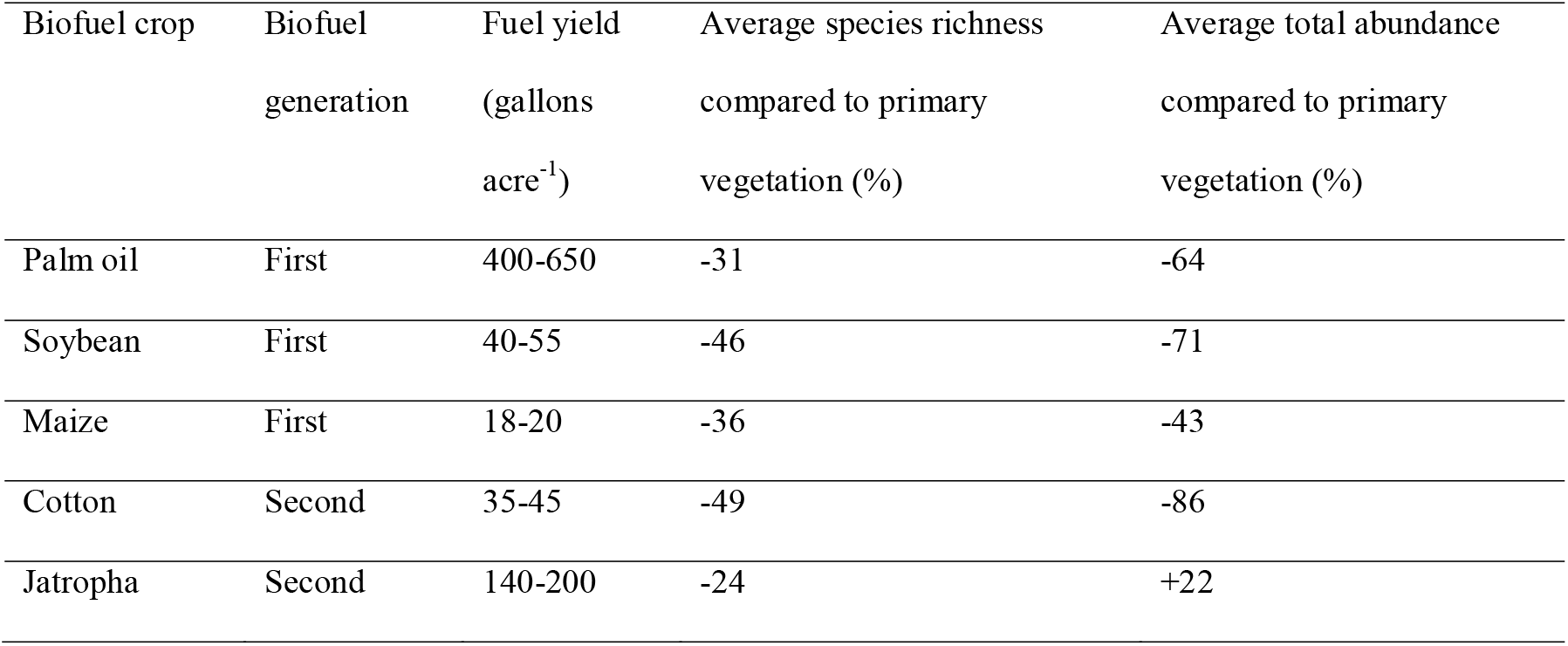
Examples of biofuel crop yield and loss of local species richness and total abundance, compared to primary vegetation. Biodiversity data are based on our analyses of data within the PREDICTS database as of March 2018. Biofuel data for jatropha are from the ‘other oil crop’ category in our analyses. Fuel yield data are based on potential fuel yields from Hoekman et al (2012).

| Biofuel crop | Biofuel generation | Fuel yield (gallons acre<sup>-1</sup>) | Average species richness compared to primary vegetation (%) | Average total abundance compared to primary vegetation (%) |
| --- | --- | --- | --- | --- |
| Palm oil | First | 400-650 | -31 | -64 |
| Soybean | First | 40-55 | -46 | -71 |
| Maize | First | 18-20 | -36 | -43 |
| Cotton | Second | 35-45 | -49 | -86 |
| Jatropha | Second | 140-200 | -24 | +22 |

### Conclusions

We have shown that biofuel crops have a negative effect on local species richness and total abundance, and that traditional first-generation biofuels are especially damaging, causing large declines in vertebrate abundance and plant species richness. Biofuels grown in Asia and Central and South America are the most detrimental, particularly oil palm and soybean, whereas in other regions there are smaller declines in biodiversity and some neutral and positive impacts. In order to minimise the destructive impacts of habitat loss on biodiversity, our results suggest that biofuel policies should not lead to further land clearance, but should instead focus on other techniques, such as using degraded land or existing waste products. Biofuel policies should be tailored to the local environment to meet both climate mitigation and biodiversity targets, with consideration given to the ecological systems in question, how they might be affected, and the yield they might produce.

