## Supplementary Material for "The impacts of biofuel crops on local biodiversity: a global synthesis"

Supplementary Material: Appendix 1

Results from our literature review, detailing the biofuel assessment of crops within the PREDICTS database as of March 2018.

| Crop name as it appears in the database | Is there evidence for biofuel potential? | Description of use for biofuel | Biofuel generation | Biofuel category | Can use waste as a biofuel? | Reference |
|---|---|---|---|---|---|---|
| *Elaeis guineensis* | Yes | African oil palm; can use oil from fruit and kernel or waste fronds, shells and trunk | First | Oil palm | Yes | Danielsen et al. (2009) Sukiran et al. (2017) |
| *Ananas comosus* | Yes | Pineapple peel | Second | Fruit/vegetable | Yes | Saladini et al. (2016) |
| *Hevea brasiliensis* | Yes | Rubber seed oil | Second | Rubber | No | Ikwuagwu et al. (2000) |
| *Annona x atemoya* | No | | | | | |
| *Dimocarpus longan* | Yes | Longan fruit tree wood can be processed to make bioethanol | Second | Fruit/vegetable | Yes | Unpaprom et al. (2019) |
| *Macadamia integrifolia* | Yes | Macadamia seed oil | First | Other oil crop | No | Rahman et al. (2016) |
| *Triticum*, Wheat, *Triticum aestivum*, *Triticum spelta* | Yes | Can use wheat grain or wheat straw, chaff, hull, husk, glumes and stems. Common wheat or spelt wheat. | First | Wheat | Yes | Tishler et al. (2015) Barman et al. (2012) Jansone and Gaile (2013) |

| | | | | | | |
|---|---|---|---|---|---|---|
| | | | | | | Godin et al. (2013) |
| *Brassica napus*, Oil seed rape | Yes | Rapeseed oil | First | Rapeseed oil | No | Malça et al. (2014) |
| *Avena sativa* | Yes | Oat grain and residue from processing- oat waste/oat straw | First | Other grain | Yes | Ahlgren et al. (2011) |
| *Solanum tuberosum* | Yes | Potato peel waste, potato pulp | Second | Fruit/vegetable | Yes | Liang and McDonald (2014)<br>Gao et al. (2012) |
| *Hordeum vulgare*, Barley | Yes | Winter barley and barley straw | First | Other grain | Yes | Ahlgren et al. (2011)<br>Nghiem et al. (2017) |
| *Sinapis alba* | Yes | Inedible seed oil from white mustard | Second | Other oil crop | No | Sáez-Bastante et al. (2016) |
| *Linum usitatissimum* | Yes | Common flax/linseed oil, seed and oil (edible) | First | Other oil crop | No | Borugadda and Goud (2012) |
| *Cucurbita pepo* | Yes | Pumpkin seed oil | First | Other oil crop | No | Schinas et al. (2009) |
| *Vicia faba* | Yes | Can use broad bean biomass residue eg. straw, can also use whole crop | First | Fruit/vegetable | Yes | Pakarinen et al. (2011)<br>Petersson et al. (2007) |

| | | | | | | |
|---|---|---|---|---|---|---|
| *Coffea arabica*, *Coffea*, Coffee | Yes | Spent coffee grounds | Second | Coffee | Yes | Gómez-de la Cruz et al. (2015) |
| *Secale cereale* | Yes | Can grow as winter crop after harvest of main summer crop, can also use rye straw | First | Other grain | Yes | Smuga-Kogut et al. (2017)<br>Shao et al. (2015) |
| Grapes | Yes | Grape skins or seeds | Second | Fruit/vegetable | Yes | Xu et al. (2009) |
| *Olea europaea* | Yes | Olive stone residue, olive pomace, olive oil extraction waste, tree pruning biomass, leaves | Second | Fruit/vegetable | Yes | Mata-Sánchez et al. (2014)<br>Dermeche et al. (2013)<br>Manzanares et al. (2017) |
| *Theobroma cacao* | Yes | Cocoa pod husk (residue after extracting pulp), cocoa pulp juice (sweatings) or cocoa pods (residue) | Second | Fruit/vegetable | Yes | Balladares et al. (2016) |
| *Medicago sativa*, Alfalfa | Yes | Can use alfalfa stems for biofuel while leaves can still be used as feed for livestock | Second | Perennial grass | Yes | Gonzalez-Garcia et al. (2010) |
| *Zea mays*, *ZeaÂ mays*, *Zea Mays*, Corn, Maize crop | Yes | Maize straw, maize silage (produced from whole plants), sugar, stover (non-grain parts; stalks, leaves and cobs remaining after harvest) and grain | First | Maize | Yes | Zbytek et al. (2016)<br>White et al. (2012)<br>Blanco-Canqui and Lal (2007) |

| | | | | | | |
|---|---|---|---|---|---|---|
| *Brassica oleracea* | No | | | | | |
| *Phaseolus vulgaris* | No | | | | | |
| *Glycine max* | Yes | Soybean oil | First | Soybean | No | Cerri et al. (2017) |
| *Gossypium* | Yes | Cotton stalk/post-harvest residue, waste cotton fabric | Second | Cotton | Yes | Christopher et al. (2017)<br>Nikolić et al. (2017) |
| *Manihot esculenta* | Yes | Cassava and cassava waste | First | Fruit/vegetable | Yes | Hanif et al. (2017)<br>Veiga et al. (2016) |
| *Colocasia esculenta* | Yes | Taro and taro waste, potential in Southeast Asia | First | Fruit/vegetable | Yes | Ben-Iwo et al. (2016) |
| *Musa x paradisiaca* | Yes | Common banana; banana lignocellulosic residue, banana peel | Second | Fruit/vegetable | Yes | Guerrero et al. (2018)<br>Oberoi et al. (2011) |
| *Solanum melongena* | No | | | | | |
| *Solanum lycopersicum* | Yes | Tomato pomace (waste skin and seeds), seed oil for biodiesel | Second | Fruit/vegetable | Yes | Allison et al. (2016) |
| *Arachis hypogaea* | Yes | Peanut; waste shells, seed oil | First | Other oil crop | Yes | Polachini et al. (2016)<br>Russo and Webber (2012) |
| *Cucumis sativus* | Yes | Cucumber pomace containing pulp, peel, seeds, and stem | Second | Fruit/vegetable | Yes | Szymanska-Chargot et al. (2017) |

| | | | | | | |
|---|---|---|---|---|---|---|
| *Daucus carota* | Yes | Wild carrot pomace, carrots which are discarded due to sizing problems | Second | Fruit/vegetable | Yes | Szymanska-Chargot et al. (2017)<br>Aimaretti et al. (2012) |
| *Apium graveolens* | No | | | | | |
| *Ribes nigrum* | Yes | Blackcurrant pomace obtained after pressing-seeds, peels and pulp | Second | Fruit/vegetable | Yes | Déniel et al. (2016) |
| *Euterpe edulis* | No | | | | | |
| *Cocos nucifera* | Yes | Coconut oil, shell, husk | First | Other oil crop | Yes | Tupufia et al. (2013)<br>Mendu et al. (2012) |
| *Psidium guajava* | No | | | | | |
| *Mangifera indica* | Yes | Mango seed oil, pulp and peel, leaf litter or stem bark residue | First | Fruit/vegetable | Yes | Akhtar et al. (2016)<br>Carrillo-Nieves et al. (2017)<br>Fernando et al. (2014) |
| *Averrhoa carambola* | No | | | | | |
| *Citrus limon* | Yes | Lemon peel waste | Second | Fruit/vegetable | Yes | Boluda-Aguilar and López-Gómez (2013) |

| | | | | | | |
|---|---|---|---|---|---|---|
| *Bambuseae* | Yes | Bamboo lignocellulosic substrate can be used due to its high growth efficiency | Second | Perennial grass | Yes | He et al. (2014) |
| *Citrus x sinensis* | Yes | Orange peel | Second | Fruit/vegetable | Yes | Negro et al. (2017) |
| *Oryza sativa* | Yes | Rice straw and husk | Second | Other grain | Yes | Banerjee et al. (2009)<br>Victor et al. (2016) |
| *Ipomoea batatas* | Yes | Starch from sweet potato tubers, residues from after separating starch, peel | First | Fruit/vegetable | Yes | Waluyo et al. 2015)<br>Wang et al. (2016) |
| *Ilex paraguariensis* | No | | | | | |
| *Avena barbata* | No | | | | | |
| *Trifolium subterraneum* | No | | | | | |
| *Helianthus annuus* | Yes | Common sunflower seed oil is commonly used in Europe, could also be used as a source of lignocellulosic biomass | First | Other oil crop | Yes | Marvey (2008)<br>Ziebell et al. (2013) |
| *Carica papaya* | Yes | Papaya peel, waste seed oil, waste fruit puree | Second | Fruit/vegetable | Yes | Dahunsi et al. (2017)<br>Winayanuwattikun et al. (2008)<br>Heller et al. (2015) |
| *Artocarpus altilis* | Yes | Breadfruit starch | First | Fruit/vegetable | No | Betiku and Taiwo (2015) |

| | | | | | | |
|---|---|---|---|---|---|---|
| *Artocarpus heterophyllus* | Yes | Jackfruit woody biomass or stone (stone is edible but most ends up as waste) | Second | Fruit/vegetable | Yes | Dutta et al. (2014)<br>Nuriana and Wuryantoro (2015) |
| *Sorghum bicolor* | Yes | Sweet sorghum stalks | First | Other grain | Yes | Chen et al. (2016)<br>Mathur et al. (2017) |
| *Eleusine coracana* | No | | | | | |
| *Panicum miliaceum* | Yes | Proso millet grain (used mainly as bird/livestock feed but can be eaten by humans). Close relative to switchgrass (well-known biofuel crop). | First | Other grain | No | Rose and Santra (2013) |
| *Pennisetum glaucum* | Yes | Ground pearl millet (grown mostly for livestock but can be eaten by humans), agricultural waste/hay | First | Other grain | Yes | Chen et al. (2007)<br>Wu et al. (2006) |
| *Setaria italica* | Yes | Foxtail millet (grown as human and animal food) | First | Other grain | No | Zhang et al. (2012) |
| Teak | Yes | Teak wood pellets | Second | Woody crop | No | Tenorio et al. (2015) |
| *Musa textilis* | No | | | | | |
| *Sechium edule* | No | | | | | |

| | | | | | | |
|---|---|---|---|---|---|---|
| *Sorghum arundinaceum* | No | | | | | |
| *Areca catechu* | Yes | Areca nut husk | Second | Woody crop | Yes | Sasmal et al. (2012) |
| *Anacardium occidentale* | Yes | Cashew nut oil, nut shell liquid (by-product from nut production), cashew apple juice (apple usually a neglected product), cashew apple bagasse (straw) | First | Other oil crop | Yes | Eddy et al. (2011)<br>Sanjeeva et al. (2014)<br>Deenanath et al. (2015)<br>Rocha et al. (2014) |
| *Inga edulis* | No | | | | | |
| *Persea americana* | Yes | Avocado flesh or seeds | First | Fruit/vegetable | Yes | Adekunle et al. (2016)<br>Aysu and Durak (2015) |
| *Camellia sinensis* | Yes | Spent kitchen waste tea can be used for biodiesel, or tea seed oil | Second | Tea | Yes | Demirbas (2010)<br>Mahmood and Hussain (2010) |
| *Ricinus communis* | Yes | Castor bean seed oil, castor plant lignocellulosic biomass- leaves and stems | Second | Other oil crop | Yes | Timko et al. (2014)<br>Mathur and Chakraborty (2016) |
| *Jatropha curcas* | Yes | Seed oil widely used as biofuel. Could also use husk/shell | Second | Other oil crop | Yes | Abhilash et al. (2011)<br>Makkar and Becker (2009) |

| | | | | | | |
|---|---|---|---|---|---|---|
| *Fragaria x ananassa* | No | | | | | |
| *Prunus persica* | Yes | Peach tree wood, bark, branches. Pruning from cultivation; discarded peaches, peels and pulp residues from processing; unsold nectar from distribution; not consumed nectar from consumption. | Second | Fruit/vegetable | Yes | Cichy et al. (2017)<br>De Menna et al. (2015)<br>Ucuncu et al. (2013) |
| *Molinia caerulea* | No | | | | | |
| *Juncus acutiï¬,orus* | No | | | | | |
| *Agrostis canina* | No | | | | | |
| *Lolium perenne* | Yes | Common ryegrass | Second | Perennial grass | | Farrar et al. (2012) |
| *Trifolium repens* | No | | | | | |
| *Protea compacta* | No | | | | | |
| *Protea cordata* | No | | | | | |
| *Leucadendron platyspermum* | No | | | | | |
| *Pisum sativum* | Yes | Pea vine waste | Second | Fruit/vegetable | Yes | Xia et al. (2016) |
| *Allium cepa* | Yes | Waste onions, juice residue, peel/skin | Second | Fruit/vegetable | Yes | Vazirzadeh et al. (2012)<br>Kim et al. (2017) |

| | | | | | | |
|---|---|---|---|---|---|---|
| *Malus domestica* | Yes | Apple seeds or pomace (waste from extracting juice) | Second | Fruit/vegetable | Yes | Górnaś and Rudzińska, (2016)<br>Gama et al. (2015) |
| *Prunus salicina* | No | | | | | |
| *Chrysanthemum cinerariaefolium* | No | | | | | |
| *Quercus suber* | No | | | | | |
| *Annona squamosa* | Yes | Custard apple seeds | Second | Other oil crop | Yes | Parthiban and Perumalsamy (2016) |
| *Bactris gasipaes* | No | | | | | |
| *Vigna unguiculata* | Yes | Cowpea biomass | Second | Fruit/vegetable | Yes | Foster et al. (2017) |
| *Eucalyptus camaldulensis* | Yes | Woody biomass | Second | Woody crop | | Acuna et al. (2017) |
| *Khaya senegalensis* | No | | | | | |
| *Dalbergia sissoo* | No | | | | | |
| *Cupressus sempervirens* | Yes | Mediterranean cypress seed oil | Second | Other oil crop | Yes | Nehdi (2013) |
| *Khaya senegalensis* | No | | | | | |

| | |
|---|---|
| *Brassica rapa* var. *rapa* | No |
| *Fagopyrum esculentum* | No |

Supplementary Material: Appendix 2

Number of sites from the PREDICTS database for each category of biofuel crop in each region and number of sites that recorded each taxonomic group for each category of biofuel crop.

| Biofuel crop category | Number of sites | | | | | | | | |
|---|---|---|---|---|---|---|---|---|---|
| | Africa | Asia | Central & South America | Europe | North America | Oceania | Invertebrates | Plants | Vertebrates |
| Coffee | 90 | 16 | 157 | 0 | 0 | 0 | 105 | 122 | 36 |
| Cotton | 35 | 0 | 0 | 0 | 0 | 0 | 24 | 11 | 0 |
| Fruit/ vegetable | 15 | 92 | 258 | 73 | 9 | 12 | 141 | 40 | 261 |
| Maize | 14 | 2 | 138 | 0 | 3 | 0 | 89 | 67 | 1 |
| Mixed crops | 682 | 43 | 272 | 94 | 71 | 33 | 175 | 338 | 682 |
| Oil palm | 0 | 74 | 0 | 0 | 0 | 0 | 28 | 0 | 46 |
| Other grain | 1 | 0 | 0 | 50 | 0 | 0 | 48 | 0 | 3 |
| Other oil crop | 4 | 6 | 1 | 4 | 22 | 9 | 29 | 6 | 9 |
| Perennial grass | 0 | 0 | 3 | 6 | 5 | 15 | 23 | 6 | 0 |
| Rapeseed oil | 0 | 1 | 0 | 79 | 0 | 9 | 89 | 0 | 0 |
| Rubber | 0 | 17 | 0 | 0 | 0 | 0 | 16 | 0 | 1 |
| Soybean | 0 | 21 | 70 | 0 | 0 | 0 | 35 | 56 | 0 |
| Wheat | 0 | 14 | 0 | 90 | 11 | 18 | 120 | 11 | 0 |

Supplementary Material: Appendix 3

Results from the model with species richness as the response variable and land-use, including biofuel crop category (LandUseCat), as the explanatory variable, with $R^2$ values and results table. SS = Source – Study, SSB = Source – Study – Block, SSBS = Source – Study - Block – Site.

Marginal $R^2$ = 0.0067 (2 sf) and conditional $R^2$ = 0.90 (2 sf). Significance of between 0 and 0.001 = ***, between 0.001 and 0.01= **, between 0.01 and 0.05 = *, between 0.05 and 0.1 = . and between 0.1 and 1 = blank space.

| Model parameter | | | | | |
|---|---|---|---|---|---|
| Random effects | Variance | SD | | | |
| SSBS | 0.07415 | 0.2723 | | | |
| SSB | 0.03362 | 0.1834 | | | |
| SS | 1.39274 | 1.1801 | | | |
| Fixed effects | Estimate | Std. Error | z value | Pr(>\|z\|) | Significance |
| (Intercept) | 2.52862 | 0.04490 | 56.32 | < 2e-16 | *** |
| LandUseCatCoffee | -0.23339 | 0.03579 | -6.52 | 6.96e-11 | *** |
| LandUseCatCotton | -0.67113 | 0.10180 | -6.59 | 4.33e-11 | *** |
| LandUseCatFruit/vegetable | -0.11646 | 0.03768 | -3.09 | 0.00200 | ** |
| LandUseCatMaize | -0.44717 | 0.04435 | -10.08 | < 2e-16 | *** |
| LandUseCatMixed crops | -0.29459 | 0.02135 | -13.80 | < 2e-16 | *** |
| LandUseCatOil palm | -0.36700 | 0.06060 | -6.06 | 1.39e-09 | *** |
| LandUseCatOther grain | -0.17718 | 0.09641 | -1.84 | 0.06610 | . |
| LandUseCatOther oil crop | -0.27909 | 0.09623 | -2.90 | 0.00373 | ** |
| LandUseCatPasture | -0.16926 | 0.01222 | -13.85 | < 2e-16 | *** |
| LandUseCatPerennial grass | -0.31261 | 0.10026 | -3.12 | 0.00182 | ** |
| LandUseCatRapeseed oil | -0.19537 | 0.07611 | -2.57 | 0.01026 | * |
| LandUseCatRubber | -0.04582 | 0.10552 | -0.43 | 0.66411 | |
| LandUseCatSecondary vegetation | -0.12534 | 0.01032 | -12.14 | < 2e-16 | *** |
| LandUseCatSoybean | -0.60632 | 0.06639 | -9.13 | < 2e-16 | *** |

| | | | | | |
|---|---|---|---|---|---|
| LandUseCatUrban | -0.25847 | 0.02368 | -10.92 | < 2e-16 | *** |
| LandUseCatWheat | -0.45508 | 0.06016 | -7.56 | 3.89e-14 | *** |

Supplementary Material: Appendix 4

Results from the model with total abundance as the response variable and land-use, including biofuel crop category (LandUseCat), as the explanatory variable, with $R^2$ values and results table. SS = Source – Study, SSB = Source – Study – Block, SSBS = Source – Study - Block – Site.

Marginal $R^2$ = 0.0033 (2 sf) and conditional $R^2$ = 0.90 (2 sf).

| Model parameter | | | |
|---|---|---|---|
| Random effects | Variance | SD | |
| SSB | 0.2240 | 0.4733 | |
| SS | 5.1455 | 2.2684 | |
| Residual | 0.6196 | 0.7871 | |
| Fixed effects | Estimate | Std. Error | t value |
| (Intercept) | 4.71044 | 0.09249 | 50.93 |
| LandUseCatCoffee | -0.32848 | 0.09917 | -3.31 |
| LandUseCatCotton | -1.99779 | 0.19839 | -10.07 |
| LandUseCatFruit/vegetable | -0.08473 | 0.07772 | -1.09 |
| LandUseCatMaize | -0.55408 | 0.10609 | -5.22 |
| LandUseCatMixed crops | -0.14496 | 0.03764 | -3.85 |
| LandUseCatOil palm | -1.01357 | 0.14475 | -7.00 |
| LandUseCatOther grain | -0.45214 | 0.14508 | -3.12 |
| LandUseCatOther oil crop | 0.20000 | 0.25439 | 0.79 |
| LandUseCatPasture | -0.21506 | 0.02556 | -8.42 |
| LandUseCatPerennial grass | 0.46546 | 0.21685 | 2.15 |
| LandUseCatRapeseed oil | 0.46843 | 0.15138 | 3.09 |
| LandUseCatRubber | -0.36355 | 0.27226 | -1.34 |
| LandUseCatSecondary vegetation | -0.17520 | 0.02219 | -7.90 |
| LandUseCatSoybean | -1.25428 | 0.26514 | -4.73 |
| LandUseCatUrban | -0.18306 | 0.04512 | -4.06 |

| | | | |
|---|---|---|---|
| LandUseCatWheat | -0.58893 | 0.10293 | -5.72 |

Supplementary Material: Appendix 5

Results from the model with total abundance as the response variable and land-use, including biofuel crop generation (LandUseGen), as the explanatory variable, with $R^2$ values and results table. SS = Source – Study, SSB = Source – Study – Block, SSBS = Source – Study - Block – Site.

Marginal $R^2$ = 0.0018 (2 sf) and conditional $R^2$ = 0.90 (2 sf).

| Model parameter | | | |
|---|---|---|---|
| Random effects | Variance | SD | |
| SSB | 0.2261 | 0.4754 | |
| SS | 5.1661 | 2.2729 | |
| Residual | 0.6225 | 0.7890 | |
| Fixed effects | Estimate | Std. Error | t value |
| (Intercept) | 4.70825 | 0.09305 | 50.60 |
| LandUseGen1st generation | -0.49234 | 0.06888 | -7.15 |
| LandUseGen2nd generation | -0.28983 | 0.05577 | -5.20 |
| LandUseGenPasture | -0.19653 | 0.02651 | -7.41 |
| LandUseGenSecondary vegetation | -0.17679 | 0.02240 | -7.89 |
| LandUseGenUrban | -0.18768 | 0.04547 | -4.13 |

Supplementary Material: Appendix 6

Results from the model with species richness as the response variable and land-use, including biofuel crop generation (LandUseGen), as the explanatory variable, with $R^2$ values and results table. SS = Source – Study, SSB = Source – Study – Block, SSBS = Source – Study - Block – Site.

Marginal $R^2$ = 0.0052 (2 sf) and conditional $R^2$ = 0.90 (2 sf). Significance of between 0 and 0.001 = ***, between 0.001 and 0.0 = **, between 0.01 and 0.05 = *, between 0.05 and 0.1= . and between 0.1 and 1 = blank space.

| Model parameter | | | | | |
|---|---|---|---|---|---|
| Random effects | Variance | SD | | | |
| SSBS | 0.07297 | 0.2701 | | | |
| SSB | 0.03276 | 0.1810 | | | |
| SS | 1.40080 | 1.1836 | | | |
| Fixed effects | Estimate | Std. Error | z value | Pr(>\|z\|) | Significance |
| (Intercept) | 2.52001 | 0.04517 | 55.79 | <2e-16 | *** |
| LandUseGen1st generation | -0.45281 | 0.02854 | -15.87 | <2e-16 | *** |
| LandUseGen2nd generation | -0.20599 | 0.02347 | -8.78 | <2e-16 | *** |
| LandUseGenPasture | -0.13778 | 0.01247 | -11.05 | <2e-16 | *** |
| LandUseGenSecondary vegetation | -0.11958 | 0.01028 | -11.63 | <2e-16 | *** |
| LandUseGenUrban | -0.25190 | 0.02362 | -10.67 | <2e-16 | *** |

Supplementary Material: Appendix 7

Results from the model with species richness as the response variable and land-use, including biofuel crop generation (LandUseGen), geographic region and their interaction as the explanatory variables, including $R^2$ values and results table.

Marginal $R^2$ = 0.022 (2 sf) and conditional $R^2$ = 0.90 (2 sf). Significance of between 0 and 0.001 = ***, between 0.001 and 0.0 = **, between 0.01 and 0.05 = *, between 0.05 and 0.1 = . and between 0.1 and 1 = blank space.

| Model parameter | | | | | |
|---|---|---|---|---|---|
| Random effects | Variance | SD | | | |
| SSBS | 0.07233 | 0.2689 | | | |
| SSB | 0.03019 | 0.1738 | | | |
| SS | 1.37324 | 1.1719 | | | |
| Fixed effects | Estimate | Std. Error | z value | Pr(>\|z\|) | Significance |
| (Intercept) | 2.390040 | 0.122233 | 19.553 | < 2e-16 | *** |
| LandUseGen1st generation | -0.444477 | 0.111503 | -3.986 | 6.71e-05 | *** |
| LandUseGen2nd generation | -0.276141 | 0.044443 | -6.213 | 5.19e-10 | *** |
| LandUseGenPasture | -0.284353 | 0.055972 | -5.080 | 3.77e-07 | *** |
| LandUseGenSecondary vegetation | -0.107792 | 0.026971 | -3.997 | 6.42e-05 | *** |
| LandUseGenUrban | -0.082235 | 0.081120 | -1.014 | 0.310702 | |
| RegionAsia | 0.402709 | 0.162717 | 2.475 | 0.013327 | * |

| | | | | | |
|---|---|---|---|---|---|
| RegionCentral & South America | 0.195139 | 0.149438 | 1.306 | 0.191616 | |
| RegionEurope | 0.047874 | 0.151953 | 0.315 | 0.752718 | |
| RegionNorth America | -0.123192 | 0.196178 | -0.628 | 0.530029 | |
| RegionOceania | 0.113362 | 0.194125 | 0.584 | 0.559245 | |
| LandUseGen1st generation:RegionAsia | -0.063970 | 0.122759 | -0.521 | 0.602293 | |
| LandUseGen2nd generation:RegionAsia | 0.127612 | 0.064390 | 1.982 | 0.047496 | * |
| LandUseGenPasture:RegionAsia | 0.069120 | 0.147680 | 0.468 | 0.639755 | |
| LandUseGenSecondary vegetation:RegionAsia | -0.098217 | 0.035725 | -2.749 | 0.005973 | ** |
| LandUseGenUrban:RegionAsia | -0.673426 | 0.133107 | -5.059 | 4.21e-07 | *** |
| LandUseGen1st generation:RegionCentral & South America | -0.027187 | 0.125005 | -0.217 | 0.827829 | |
| LandUseGen2nd generation:RegionCentral & South America | 0.051698 | 0.061567 | 0.840 | 0.401078 | |
| LandUseGenPasture:RegionCentral & South America | 0.163596 | 0.060684 | 2.696 | 0.007021 | ** |
| LandUseGenSecondary vegetation:RegionCentral & South America | 0.162352 | 0.034389 | 4.721 | 2.35e-06 | *** |
| LandUseGenUrban:RegionCentral & South America | -0.416793 | 0.166678 | -2.501 | 0.012399 | * |
| LandUseGen1st generation:RegionEurope | 0.075298 | 0.124818 | 0.603 | 0.546334 | |
| LandUseGen2nd generation:RegionEurope | 0.230661 | 0.083655 | 2.757 | 0.005828 | ** |
| LandUseGenPasture:RegionEurope | 0.157195 | 0.061616 | 2.551 | 0.010735 | * |
| LandUseGenSecondary vegetation:RegionEurope | -0.128369 | 0.036306 | -3.536 | 0.000407 | *** |

| | | | | | |
|---|---|---|---|---|---|
| LandUseGenUrban:RegionEurope | -0.223665 | 0.088725 | -2.521 | 0.011706 | * |
| LandUseGen1st generation:RegionNorth America | 0.048959 | 0.278482 | 0.176 | 0.860447 | |
| LandUseGen2nd generation:RegionNorth America | 0.006152 | 0.192933 | 0.032 | 0.974562 | |
| LandUseGenPasture:RegionNorth America | -0.075363 | 0.086385 | -0.872 | 0.382982 | |
| LandUseGenSecondary vegetation:RegionNorth America | 0.102169 | 0.043984 | 2.323 | 0.020185 | * |
| LandUseGenUrban:RegionNorth America | -0.101399 | 0.093512 | -1.084 | 0.278215 | |
| LandUseGen1st generation:RegionOceania | 0.071064 | 0.154608 | 0.460 | 0.645774 | |
| LandUseGen2nd generation:RegionOceania | 0.068285 | 0.136288 | 0.501 | 0.616348 | |
| LandUseGenPasture:RegionOceania | 0.099361 | 0.061165 | 1.624 | 0.104277 | |
| LandUseGenSecondary vegetation:RegionOceania | -0.002824 | 0.043552 | -0.065 | 0.948307 | |
| LandUseGenUrban:RegionOceania | 0.045525 | 0.161379 | 0.282 | 0.777866 | |

Supplementary Material: Appendix 8

Results from the model with total abundance as the response variable and land-use, including biofuel crop generation (LandUseGen), geographic region and their interaction as the explanatory variables, including $R^2$ values and results table.

Marginal $R^2$ = 0.022 (2 sf) and conditional $R^2$ = 0.90 (2 sf).

| Model parameter | | | |
|---|---|---|---|
| Random effects | Variance | SD | |
| SSB | 0.2154 | 0.4641 | |
| SS | 5.1612 | 2.2718 | |
| Residual | 0.6131 | 0.7830 | |
| Fixed effects | Estimate | Std. Error | t value |
| (Intercept) | 4.14540 | 0.26010 | 15.938 |
| LandUseGen1st generation | -0.44274 | 0.30092 | -1.471 |
| LandUseGen2nd generation | -0.90400 | 0.15224 | -5.938 |
| LandUseGenPasture | -0.39018 | 0.08931 | -4.369 |
| LandUseGenSecondary vegetation | -0.10541 | 0.05674 | -1.858 |
| LandUseGenUrban | 0.65901 | 0.12814 | 5.143 |
| RegionAsia | 0.64724 | 0.34881 | 1.856 |
| RegionCentral & South America | 0.55443 | 0.31337 | 1.769 |
| RegionEurope | 0.93837 | 0.32114 | 2.922 |
| RegionNorth America | 0.09253 | 0.41082 | 0.225 |
| RegionOceania | 1.26998 | 0.39832 | 3.188 |
| LandUseGen1st generation:RegionAsia | -0.62938 | 0.32615 | -1.930 |
| LandUseGen2nd generation:RegionAsia | 0.46405 | 0.19940 | 2.327 |
| LandUseGenPasture:RegionAsia | 0.08827 | 0.36299 | 0.243 |
| LandUseGenSecondary vegetation:RegionAsia | -0.10407 | 0.08374 | -1.243 |
| LandUseGenUrban:RegionAsia | -1.71004 | 0.27131 | -6.303 |

| | | | |
|---|---|---|---|
| LandUseGen1st generation:RegionCentral & South America | -0.67156 | 0.37873 | -1.773 |
| LandUseGen2nd generation:RegionCentral & South America | 0.60569 | 0.17723 | 3.417 |
| LandUseGenPasture:RegionCentral & South America | 0.34997 | 0.10020 | 3.493 |
| LandUseGenSecondary vegetation:RegionCentral & South America | 0.21553 | 0.07110 | 3.031 |
| LandUseGenUrban:RegionCentral & South America | -1.49846 | 0.26502 | -5.654 |
| LandUseGen1st generation:RegionEurope | -0.14400 | 0.32161 | -0.448 |
| LandUseGen2nd generation:RegionEurope | 0.53359 | 0.19448 | 2.744 |
| LandUseGenPasture:RegionEurope | -0.07174 | 0.10443 | -0.687 |
| LandUseGenSecondary vegetation:RegionEurope | -0.56198 | 0.07691 | -7.307 |
| LandUseGenUrban:RegionEurope | -1.28504 | 0.14921 | -8.612 |
| LandUseGen1st generation:RegionNorth America | 0.30402 | 0.61347 | 0.496 |
| LandUseGen2nd generation:RegionNorth America | 2.12108 | 0.38889 | 5.454 |
| LandUseGenPasture:RegionNorth America | -0.10123 | 0.14481 | -0.699 |
| LandUseGenSecondary vegetation:RegionNorth America | 0.31089 | 0.08956 | 3.471 |
| LandUseGenUrban:RegionNorth America | -0.74287 | 0.15427 | -4.816 |
| LandUseGen1st generation:RegionOceania | 0.37073 | 0.35965 | 1.031 |
| LandUseGen2nd generation:RegionOceania | 1.07214 | 0.28005 | 3.828 |
| LandUseGenPasture:RegionOceania | -0.01691 | 0.11180 | -0.151 |
| LandUseGenSecondary vegetation:RegionOceania | -0.13357 | 0.09171 | -1.456 |
| LandUseGenUrban:RegionOceania | -0.95576 | 0.25668 | -3.724 |

Supplementary Material: Appendix 9

Results from the model with species richness as the response variable and land-use, including biofuel crop generation (LandUseGen), taxon and their interaction as the explanatory variables, including $R^2$ values and results table.

Marginal $R^2$ = 0.050 (2 sf) and conditional $R^2$ = 0.91 (2 sf). Significance of between 0 and 0.001 = ***, between 0.001 and 0.0 = **, between 0.01 and 0.05 = *, between 0.05 and 0.1 = . and between 0.1 and 1 = blank space.

| Model parameter | | | | | |
|---|---|---|---|---|---|
| Random effects | Variance | SD | | | |
| SSBS | 0.06778 | 0.2604 | | | |
| SSB | 0.03364 | 0.1834 | | | |
| SS | 1.33408 | 1.1550 | | | |
| Fixed effects | Estimate | Std. Error | z value | Pr(>\|z\|) | Significance |
| (Intercept) | 2.55312 | 0.06127 | 41.67 | < 2e-16 | *** |
| LandUseGen1st generation | -0.37451 | 0.03973 | -9.43 | < 2e-16 | *** |
| LandUseGen2nd generation | -0.19710 | 0.04093 | -4.82 | 1.47e-06 | *** |
| LandUseGenPasture | -0.14665 | 0.01894 | -7.74 | 9.70e-15 | *** |
| LandUseGenSecondary vegetation | -0.08747 | 0.01653 | -5.29 | 1.22e-07 | *** |
| LandUseGenUrban | -0.17225 | 0.03147 | -5.47 | 4.40e-08 | *** |
| TaxonPlants | 0.37441 | 0.12186 | 3.07 | 0.002123 | ** |
| TaxonVertebrates | -0.35547 | 0.10283 | -3.46 | 0.000547 | *** |
| LandUseGen1st generation:TaxonPlants | -0.30122 | 0.06363 | -4.73 | 2.20e-06 | *** |
| LandUseGen2nd generation:TaxonPlants | -0.12026 | 0.05586 | -2.15 | 0.031322 | * |
| LandUseGenPasture:TaxonPlants | 0.12277 | 0.02781 | 4.41 | 1.01e-05 | *** |
| LandUseGenSecondary vegetation:TaxonPlants | -0.13365 | 0.02443 | -5.47 | 4.47e-08 | *** |
| LandUseGenUrban:TaxonPlants | -0.13112 | 0.06294 | -2.08 | 0.037227 | * |

| | | | | | |
|---|---|---|---|---|---|
| LandUseGen1st generation:TaxonVertebrates | 0.03905 | 0.08070 | 0.48 | 0.628425 | |
| LandUseGen2nd generation:TaxonVertebrates | 0.02807 | 0.06088 | 0.46 | 0.644774 | |
| LandUseGenPasture:TaxonVertebrates | -0.20913 | 0.03626 | -5.77 | 8.06e-09 | *** |
| LandUseGenSecondary vegetation:TaxonVertebrates | -0.01520 | 0.02593 | -0.59 | 0.557703 | |
| LandUseGenUrban:TaxonVertebrates | 0.04899 | 0.06907 | 0.71 | 0.478175 | |

Supplementary Material: Appendix 10

Results from the model with total abundance as the response variable and land-use, including biofuel crop generation (LandUseGen), taxon and their interaction as the explanatory variables, including $R^2$ values and results table.

Marginal $R^2$ = 0.066 (2 sf) and conditional $R^2$ = 0.90 (2 sf).

| Model parameter | | | |
|---|---|---|---|
| Random effects | Variance | SD | |
| SSB | 0.2285 | 0.4780 | |
| SS | 4.8461 | 2.2014 | |
| Residual | 0.6120 | 0.7823 | |
| Fixed effects | Estimate | Std. Error | t value |
| (Intercept) | 5.03754 | 0.12236 | 41.17 |
| LandUseGen1st generation | -0.36756 | 0.07929 | -4.64 |
| LandUseGen2nd generation | -0.27573 | 0.08167 | -3.38 |
| LandUseGenPasture | -0.03758 | 0.03819 | -0.98 |
| LandUseGenSecondary vegetation | -0.09210 | 0.03395 | -2.71 |
| LandUseGenUrban | -0.09720 | 0.05646 | -1.72 |
| TaxonPlants | 0.15314 | 0.26414 | 0.58 |
| TaxonVertebrates | -1.32267 | 0.20717 | -6.38 |
| LandUseGen1st generation:TaxonPlants | 0.02696 | 0.24243 | 0.11 |
| LandUseGen2nd generation:TaxonPlants | -0.33127 | 0.15823 | -2.09 |
| LandUseGenPasture:TaxonPlants | -0.18037 | 0.06492 | -2.78 |
| LandUseGenSecondary vegetation:TaxonPlants | -0.39992 | 0.05675 | -7.05 |
| LandUseGenUrban:TaxonPlants | -0.26827 | 0.14549 | -1.84 |
| LandUseGen1st generation:TaxonVertebrates | -0.77699 | 0.20997 | -3.70 |
| LandUseGen2nd generation:TaxonVertebrates | 0.04455 | 0.12921 | 0.34 |
| LandUseGenPasture:TaxonVertebrates | -0.48654 | 0.06721 | -7.24 |

| | | | |
|---|---|---|---|
| LandUseGenSecondary vegetation:TaxonVertebrates | 0.00707 | 0.05351 | 0.13 |
| LandUseGenUrban:TaxonVertebrates | 0.45410 | 0.13006 | 3.49 |
